# Machine learning and multi-omic analysis reveal contrasting recombination landscape of A and C subgenomes of winter oilseed rape

**DOI:** 10.1101/2025.09.23.677995

**Authors:** Jose A. Montero-Tena, Silvia F. Zanini, Gözde Yildiz, Tobias Kox, Amine Abbadi, Rod J. Snowdon, Agnieszka A. Golicz

## Abstract

Meiotic recombination is essential for generating genetic diversity, driving plant evolution, and enabling crop improvement, yet its uneven distribution across genomes constrains breeding efforts. Here, we investigated the multi-omic landmarks that shape the recombination landscape in *Brassica napus* by integrating epigenomic, genomic and transcriptomic data with recombination maps derived from large multi-parental rapeseed populations. Predictive machine-learning accurately predicted recombination rates and hotspot location using only feature information. Recombination was generally suppressed in centromeres and other repeat-rich, methylated regions and enriched in gene-dense, transcriptionally active domains. Proxies for chromatin configuration—such as DNA methylation, transposable elements or genes— consistently achieved the highest predictive power with the random forest algorithm. We discovered distinct recombination landscape patterns between subgenomes, with crossovers clustering near subtelomeres in the A subgenome and more evenly spread across the C subgenome. Models trained on A-subgenome data outperformed those based on the C subgenome, although combining both subgenomes improved overall accuracy.

## Introduction

Meiotic recombination is the exchange of DNA between homologous chromosomes. This highly conserved mechanism is observed universally across eukaryotes (Bernstein and Bernstein, 2010) and ensures accurate segregation of chromosomes during meiosis, maintaining genomic integrity (Lambing *et al*., 2017). In addition, meiotic recombination promotes genetic diversity by generating new allele combinations, playing a central role in plant evolution and crop improvement (Wijnker and Jong, 2008).

*Brassica napus* (oilseed rape; rapeseed) is one of the youngest cultivated crop species and one of the most important oilseed crops worldwide. It is an allopolyploid species (AACC, 2n = 38) that originated from the interspecific hybridization of *Brassica rapa* (A subgenome) and *Brassica oleracea* (C subgenome), which caused a genetic bottleneck (Allender and King, 2010). Genetic diversity was further eroded by strong selection, including extreme selection for essential seed quality traits, such as oil yield or oil quality (Qian *et al*., 2014). To generate new genetic diversity, it is essential to harness the potential of meiotic recombination, requiring a deeper understanding of the factors underlying the frequency and positioning of meiotic recombination events in rapeseed.

Across plant genomes, recombination is not uniformly distributed. A near-universal feature is the suppression of crossovers (COs) in centromeric regions, where recombination is constrained to ensure proper chromosome segregation (Vincenten *et al*., 2015). Instead, crossovers tend to cluster in recombination “hotspots” (Lambing *et al*., 2017); rapeseed exemplifies this pattern, with COs occurring in only 38% of its genome (Boideau *et al*., 2022). In maize, recombination is largely absent from pericentromeric regions but increases toward subtelomeric areas before decreasing at telomeric ends (Rodgers Melnick *et al*., 2015). However, this spatial pattern is not conserved across all plant species; in *Arabidopsis* and rice, for instance, recombination is distributed more broadly across non-centromeric regions (Salomé *et al*., 2012; Si *et al*., 2015). Across plant genomes, recombination is suppressed in heavily methylated, rich in transposable elements (TEs), heterochromatic regions but favored in low-methylation, TE-poor, euchromatic regions with open chromatin, highlighting a conserved inverse relationship between crossover frequency and both DNA methylation and TE density (Rodgers Melnick *et al*., 2015; Wijnker *et al*., 2013). In rapeseed, the same antagonism is seen in the distal ends, where large non-recombinant “islands” colocalize with elevated DNA methylation (Boideau *et al*., 2022). These islands reinforce the idea that methylation-mediated silencing of mutagenic TEs is a well-conserved, principal force suppressing crossovers throughout the genome (Yelina *et al*., 2015).

Not all methylation contexts affect recombination equally. Whereas symmetrical, heritable (Law and Jacobsen, 2010) CpG and CHG methylation is typically anti-recombinogenic, asymmetric CHH methylation shows a positive—albeit variable—association with crossover frequency in rice (Peñuela *et al*., 2022), tomato, sorghum, *Arabidopsis* (Peñuela *et al*., 2024), and maize (Peñuela *et al*., 2022; Rodgers Melnick *et al*., 2015). This positive link may reflect its role suppressing mutagenic TEs near genes, as reported in maize (Gent *et al*., 2013). Although genes have frequently been associated with elevated recombination rates, their predictive value appears to vary among plant species. In maize and tomato, crossover regions of 10 Kbp were enriched in gene-associated features, whereas in rice and *Arabidopsis*, such regions contained fewer genes (Demirci *et al*., 2018). Low-recombination regions are also associated with higher genetic load, likely due to reduced efficiency of selective sweeps and limited removal of deleterious mutations (Rodgers Melnick *et al*., 2015).

Nucleotide composition has also been implicated in shaping recombination landscapes. In maize, GC content shows a strong positive correlation with crossover frequency, possibly due to GC-biased gene conversion (Rodgers Melnick *et al*., 2015). Conversely, in *Arabidopsis*, crossover hotspots tend to be GC-poor and are enriched for AT/TA dinucleotides—associated with gene promotors and open chromatin states—which also serve as predictors of crossover regions in tomato (Demirci *et al*., 2018; Wijnker *et al*., 2013).

In rapeseed, the A and C subgenomes may differ in crossover activity. The A subgenome exhibits a higher mean crossover frequency than the C subgenome (Yan *et al*., 2023), in parallel with its enrichment for active chromatin marks that promote CO formation (Zhang *et al*., 2021). Whether the genome-wide recombination pattern in rapeseed is diploid-like, uniform across subgenomes, or subgenome-specific remains unresolved.

In this study, we aim to characterize the genome-wide recombination landscape of rapeseed. We built recombination maps from detected meiotic recombination events based on SNP data from large multi-parental populations of German winter oilseed rape. We integrated recombination data with multi-omics data consisting in various features associated with recombination— including DNA methylation, annotated sequences or telomeric proximity. The power of these features for predicting recombination, as well as the interactions between them, were quantified using a machine learning approach. We confirmed some of the well-conserved trends established previously that shape recombination genome-wide, explored the contrast in pro- and anti-recombinatory features between the A and C subgenome, and dissected the particularities of TE body CHH methylation in rapeseed. Interestingly, we identified differences in the recombination landscape between the A and C subgenome, with crossovers occurring predominantly in subtelomeric regions of the A subgenome, whereas they are more frequent in pericentromeric regions of the C subgenome.

## Results

### Generation of a high-stringent recombination data set

The raw set of 171,276 crossovers were detected across 5,132 meiotic events. Because of the very high number of individuals to be genotyped, we used a 15K single nucleotide polymorphism (SNP) genotyping array that offered moderate resolution, with a median interval length of 0.87 Mbp and a mean of 2.63 Mbp (See Supplementary Figure S1).

To improve data quality we applied two stringent filters. First, plants carrying more than 100 COs—approximately 2.6 COs per chromosome per meiosis, well above the expected 1.2 CO per chromosome (Yan *et al*., 2023)—were removed as likely genotyping artefacts (See Supplementary Figure S3). Second, COs whose intervals exceeded 2 Mbp were discarded to eliminate the long tail in the length distribution. After removing ∼13% of the original COs, these steps increased resolution (median length 0.45 Mbp, mean length 0.613 Mbp), and strengthened correlations among population-wide recombination profiles (See Supplementary Figure S4). The resulting landscapes, built with 0.3-Mbp resolution, closely resembled those reported in earlier rapeseed maps (Yan *et al*., 2023), confirming the reliability of the filtered dataset.

### Features shape the genome-wide recombination landscape

We observed strong genome-wide associations between CO formation and epigenomic, genomic, and transcriptomic features (See Figure 1). CpG methylation levels exhibited a strong positive correlation with transposable elements and an inverse relationship with gene density and transcriptional activity. In rapeseed, the recombination landscape was markedly biased, with recombination rates negatively correlated with methylated, TE-dense genomic regions, and positively correlated with gene-rich, actively transcribed regions. COs were largely suppressed in centromeric regions, characterized by “anti-recombinatory” feature profiles—high DNA methylation, dense TE accumulation, low gene content, and low gene expression. In contrast, recombination frequency increased progressively toward the telomeric regions, paralleling pronounced “pro-recombinatory” feature profiles, and peaked near the chromosome ends before declining.

**Figure 1.**
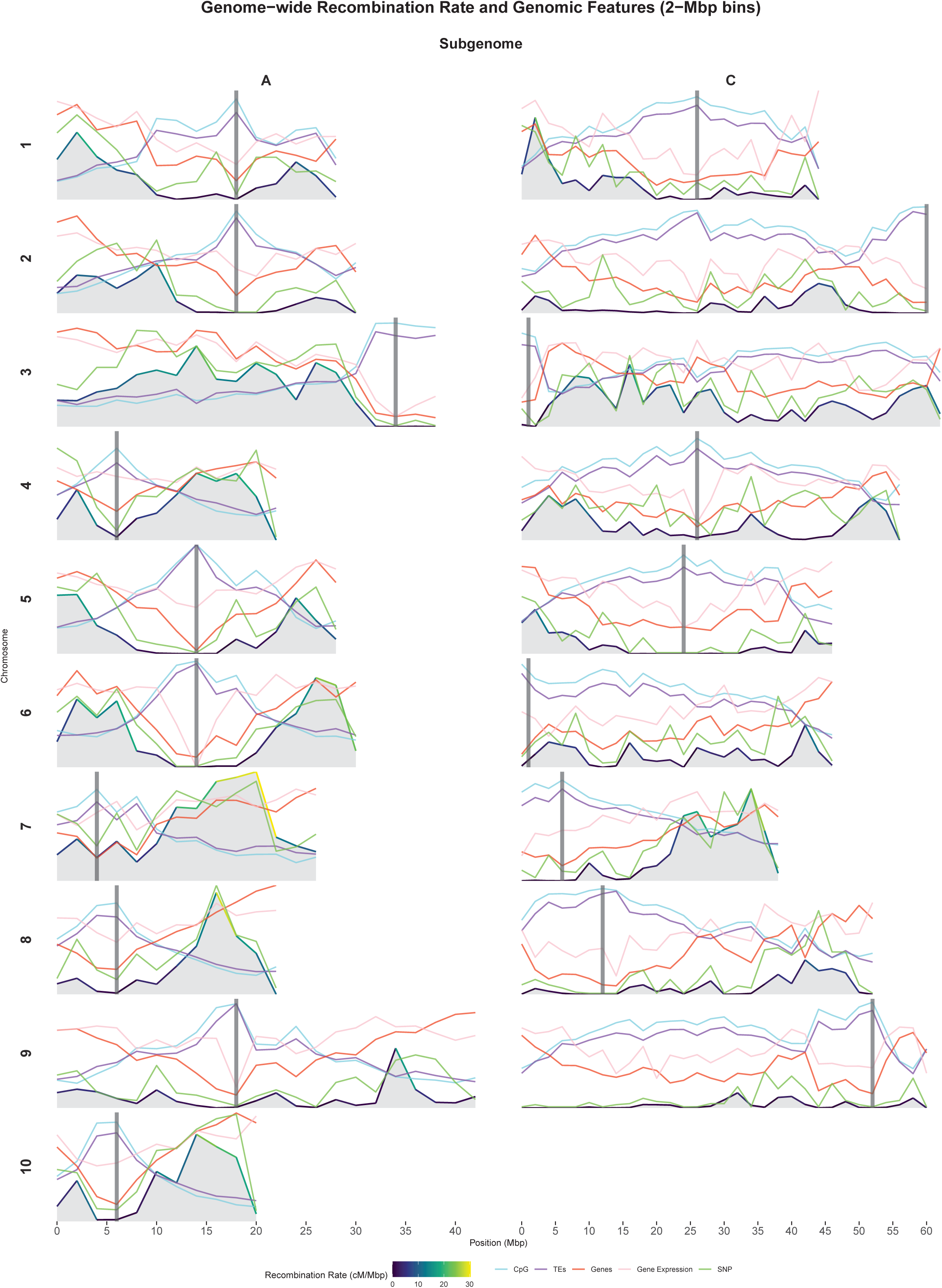
Genome-wide recombination rate and feature values over 2-Mbp genomic bins. The grey areas account for recombination rate, in cM/Mbp, as well as the color of the overlying line representing its value. The features, CpG methylation rate, TE fraction, gene fraction, gene expression level (log_10_ TPM + 0.1), SNP content, are normalized by the genome-wide maximum, and represented by light blue, purple, red, pink and green lines, respectively. Chromosomes are sorted column-wise by subgenome and row-wise by number. Centromere locations are indicated with thick dark grey vertical lines.

We observed suppression of recombination near the centromeres uniformly across chromosomes, with additional declines near telomeres. In the A subgenome, crossovers seem to modestly concentrate in subtelomeric euchromatin, whereas in the C subgenome they are distributed more irregularly along the arms, yielding a mosaic landscape with no clear dominant zone. These depressions in recombination rate across the chromosome arms resemble the dips reported in the maize genome, and might harbor gene-rich regions where low CO frequency limits selective sweeps, contributing to the retention of deleterious alleles (Rodgers Melnick *et al*., 2015).

The genome-wide mean and median recombination rates were 5.271 and 2.32 centimorgan per megabase pair (cM/Mb), respectively. However, estimating recombination rate precisely in large populations is a complicated task, with different strategies of dealing with false positive COs (Montero-Tena *et al*., 2024), which can lead to some discrepancies in estimates (Wang *et al*., 2023; Yan *et al*., 2023).

Crossover estimates are influenced by the uneven SNP density of the 15 K array. SNP-rich windows inflate apparent recombination rates, whereas marker-poor windows appear as recombination deserts. Normalizing CO counts by SNPs per bin preserved the genome-wide pattern (see Supplementary Figure S6), agreeing with the positive association between genetic polymorphisms and recombination (Cutter and Payseur, 2013; Hsu *et al*., 2022; Yang *et al*., 2015). We therefore excluded bins with no markers, as their zero CO rate are likely artefacts. For example, the distal end of chrC09L lacks array markers yet contains genes and shows CO activity in other studies (Yan *et al*., 2023). Although removing such “SNP deserts” lowers spatial resolution and may hide genuine low-recombination domains, it yields a dataset that can be compared fairly across bins.

The highest 5% of bins displayed recombination rates at least four times greater than the genome-wide mean, highlighting pronounced hotspots within the recombination landscape (See Figure 2). The *t*-test confirmed the expected antagonism between pro- and anti-recombinatory features in hotspots (See Supplementary Figure S5). Hotspots showed significantly less DNA methylation in the CpG, CHG, and CHH contexts, and lower transposon/retrotransposon coverage than randomly sampled regions. Conversely, gene expression and gene density were higher in hotspots, and chromatin accessibility also trended upward (*P* > 0.1, not significant). Additionally, the lower values of the distance between genomic bins and the closest telomere in hotspots indicate that hotspots lay closer to telomeres, whereas non-hotspots were drawn mainly from more centromeric regions. Together, these contrasts support a model in which COs concentrated in hypomethylated, gene-rich, TE-poor euchromatic domains with accessible chromatin (Rodgers Melnick *et al*., 2015; Yelina *et al*., 2012, 2015).

**Figure 2.**
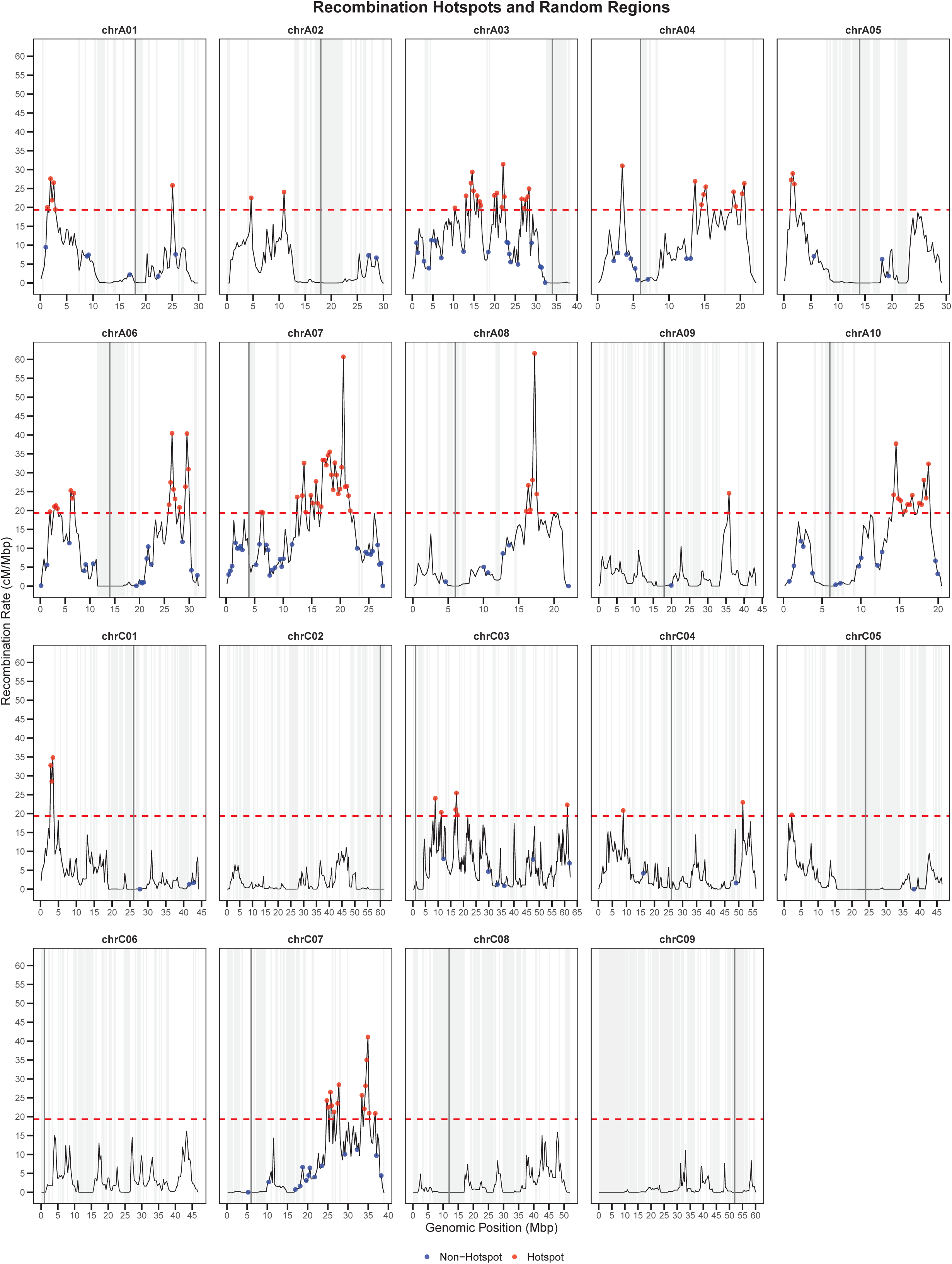
Genome-wide distribution of hotspots and non-hotspots regions. Hotspots (red dots) were in the upper 5% genome-wide recombination rate, whereas non-hotspots (blue dots) were randomly drafted from the bottom 75%, excluding SNP-empty bins. Chromosomes are labeled over the plot frames. Centromere locations are indicated with thick dark grey vertical lines. SNP-emtpy bins are colored in light grey.

### Subgenome-based differences in recombination and multi-omic features

The A and C subgenomes exhibited distinct crossover landscapes, with recombination rates that were higher in A (mean = 8.00 cM/Mbp; median = 5.31 cM/Mbp) than in C (mean = 3.54 cM/Mbp; median = 1.44 cM/Mbp), confirming earlier findings (Wang *et al*., 2023; Yan *et al*., 2023). Despite its larger physical size, the C subgenome yielded a similar absolute number of COs to the A subgenome (See Supplementary Figure S7). This discrepancy is largely attributable to its lower SNP-marker density (See Supplementary Figure S8)—in line with the lower genetic diversity of the C subgenome (Qian *et al*., 2014)—, which limits breakpoint resolution. Because recombination rarely reached the genome-wide hotspot threshold in the C subgenome, only a few of its bins qualified—none on chromosomes C02, C06, C08, or C09—whereas hotspots were far more common in the A subgenome (See Figure 2). Furthermore, the C subgenome contained broader non-recombinant regions that extend beyond centromeres (See Figure 1).

Beyond differences in CO frequency and marker density, the two subgenomes diverged sharply in the chromatin features linked to crossover formation (See Figure 3; See Supplementary Figure S10). The C subgenome carried higher DNA methylation in the CpG, CHG and CHH contexts, and greater transposable-element coverage—features associated with recombination suppression—while the A subgenome was richer in genes and transcripts. Our observations agreed with previous findings (Wang *et al*., 2018). These disparities generated pronounced bimodal distributions, especially for DNA methylation and gene density (See Figure 3; See Supplementary Figure S10): most C-subgenome bins clustered in a high-methylation, low-gene mode, whereas A-subgenome bins occupied a low-methylation, high-gene mode. These contrasts parallel the asymmetric epigenome reported by Zhang et al.: the A subgenome is enriched for active histone marks and shows higher gene content and transcriptional output, whereas the C subgenome is more heavily methylated, TE-rich, and transcriptionally subdued (Zhang *et al*., 2021).

**Figure 3.**
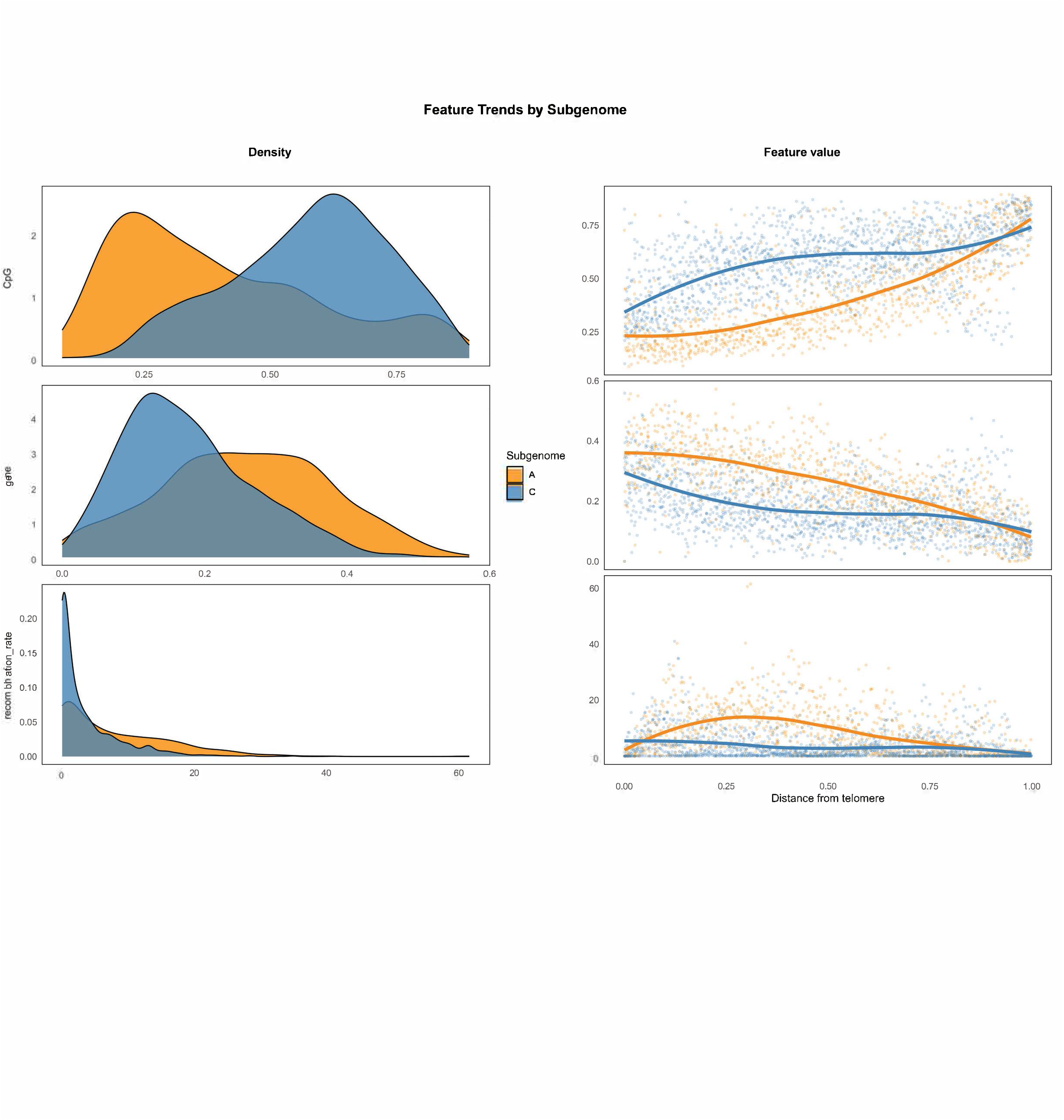
Density curve (left) and distribution across the chromosome arms (right) of feature values in the A (orange) and C (blue) subgenome for CpG methylation rate (top), gene coverage (middle) and recombination rate (bottom). The feature ‘distance from the telomere’ is represented in the X axis of the scatterplot. LOESS regression lines were fit to the scatterplot data points of the A (orange) and C (blue) subgenomes. See Supplementary Figure S10 for extended version including more features.

### Positive association of TE body CHH methylation rate and recombination

Across the genome, CHH methylation tends to occur in recombination-poor regions, similar to CpG and CHG methylation (See Supplementary Figure S11). However, when CHH sites were restricted to transposable element (TE) bodies, methylation was positively correlated with local CO rates and its levels were higher (∼8–14%) than for CHH sites across all genomic contexts (∼0–7%), in which it was, instead, negatively correlated (see Figure 4). Despite the overall higher CHH methylation levels in the C subgenome, TE bodies exhibited slightly higher CHH methylation in the A subgenome, consistent with its enrichment in pro-recombinatory features (See Supplementary Figure S10). This agrees with findings in rice, sorghum, tomato, and maize (Peñuela *et al*., 2022, 2024; Rodgers Melnick *et al*., 2015), where a positive association was reported even without restricting CHH methylation to TE bodies.

**Figure 4.**
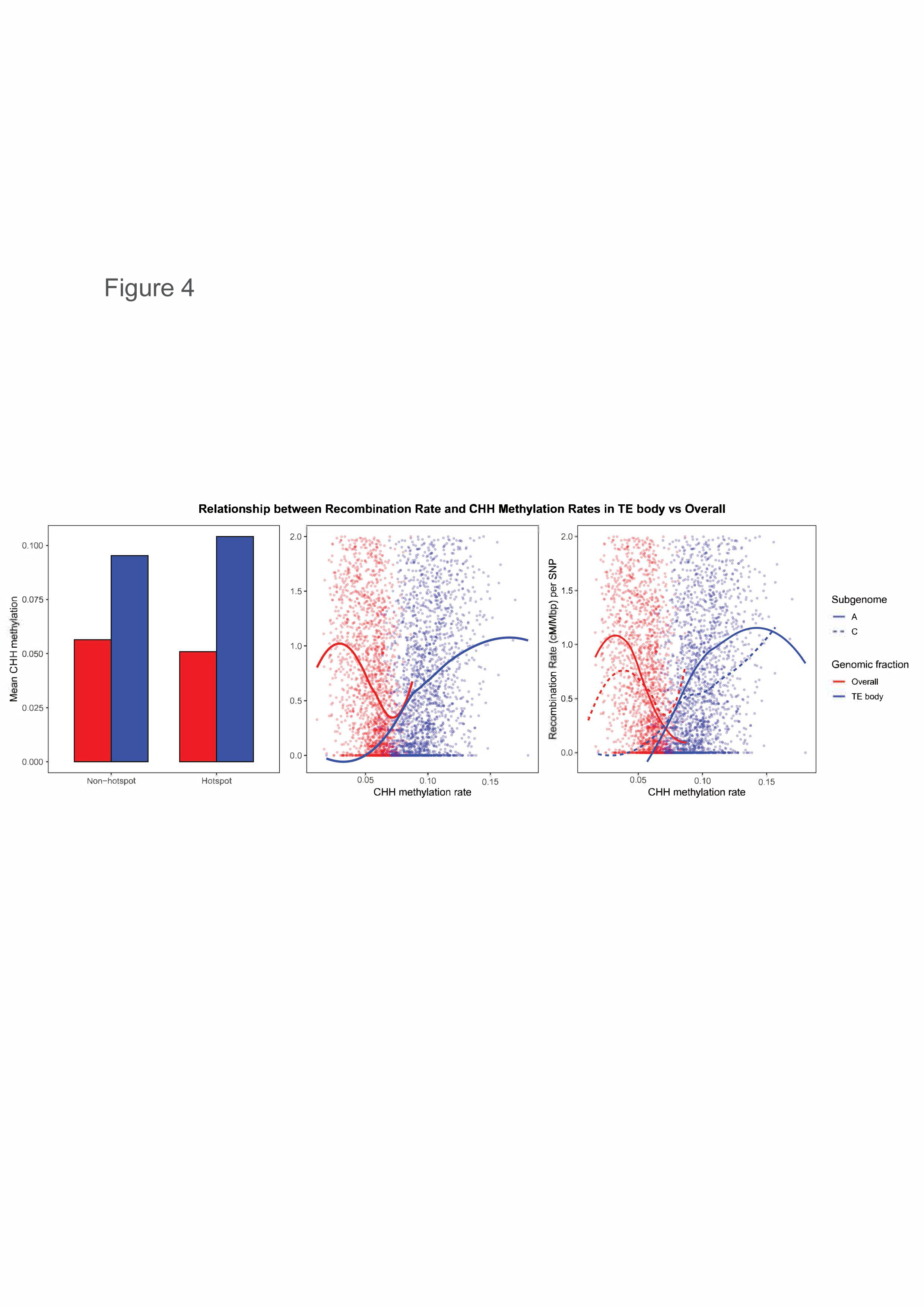
CHH methylation rate by genomic context in hotspots and non-hotspot bins (left) and relationship between CHH methylation rate and recombination rate: Genome-wide (middle), and by subgenomes (left). Two genomic contexts are compared: All CHH’s regardless of the context (overall, blue) and CHH’s exclusively located within TE bodies (TE body, red). Recombination rate is measured in cM/Mbp. LOESS regression lines were fit to the scatterplots, whereby the straight lines correspond with the A subgenome and the dashed lines correspond with the C subgenome in the left-most figure.

Although CHH methylation represents a relatively small proportion of total methylation (Law and Jacobsen, 2010), it plays a key role in complex regulatory mechanisms that suppress transposable elements (Stroud *et al*., 2014). In maize, TEs inserted upstream of genes—often in recombination-rich regions—are transcriptionally silenced via RNA-dependent CHH methylation, forming “CHH islands” (Gent *et al*., 2013). In Arabidopsis, region-specific CHH methylation pathways have been described: DRM2 silences short TEs at distal chromosomal ends, while CMT2 targets long, pericentromeric TEs, both acting in self-reinforcing loops with histone modifications and small interfering RNAs (siRNA; Stroud *et al*., 2014). Moreover, CO relocations toward distal ends have been observed in mutants lacking MOP1, a key component of the RNA-dependent CHH methylation pathway (Zhao *et al*., 2021).

Our results confirm the positive association of recombination and CHH methylation within TE bodies in rapeseed, which also display pronounced hypermethylation compared to other genome fractions. Additionally, positive correlations are observed for TE-body CHH methylation with genes and gene expression (See Supplementary Figure S11). These observations support the idea that, inside recombination hotspots, CHH methylation marks gene-proximal TE bodies strongly, preventing their activation via RNA polymerase–mediated transcription. This co-localization of TE-body CHH methylation with crossover (CO) sites does not necessarily indicate that CHH methylation promotes recombination, as contrasting patterns have been reported in *Arabidopsis* and maize CHH methylation mutants (Christophorou *et al*., 2020; Zhao *et al*., 2021). CHH methylation within TE bodies negatively correlates with TE coverage per bin, especially in retrotransposons (see Supplementary Figure S11), which is consistent with the finding that most gene-proximal LTR retrotransposons are low-copy (Grandbastien, 2015). However, the contrasting context-specific (TE vs all genomic contexts) observations could be explained by the pronounced separation between genes and TEs that we observed in the rapeseed genome (see Supplementary Figure S11), which could affect the CHH methylation signal at the resolution applied in this study. In *Arabidopsis*, a similarly contrasting distribution between TEs and genes (Zhang, 2008) could explain the weak overall positive association between CHH methylation and CO frequency (Peñuela *et al*., 2024). Overall, TE-body methylation of CHH islands and its association with recombination are complex biological processes, which could follow species-dependent trends (Martin *et al*., 2021).

### Machine learning with genome-wide data

We implemented four machine learning algorithms—regularized linear/logistic regression (LR) and three tree-based models: decision tree (DT), random forest (RF), and gradient boosting (GB)—to identify multi-omics predictors of recombination. Models were applied to two tasks: (i) classification, to distinguish recombination hotspots from non-hotspot regions, and (ii) regression, to predict recombination rate per genomic bin. These models were used to evaluate the predictive value of individual features and to assess consistency in feature importance across tasks and algorithms.

### Model sensitivity to multicollinearity in feature importance estimation

The recombination landscape in rapeseed is controlled by highly correlated features. As our goal was to evaluate the predictive value of recombination rate and hotspot distribution among these features, presumed to contribute similarly to the model’s predictions, we inspected the robustness of different machine learning models under clusters of co-linear features (see Materials and methods). Among all models tested, RF consistently produced the most stable rankings of feature importance, maintaining over 97% Spearman correlation between rankings obtained with and without highly correlated features (See Table 1; See Supplementary Figure S11). RF exhibited the lowest intra-cluster variability in importance scores among correlated features, indicating a more balanced attribution of predictive power across co-linear variables. Additionally, the correlation in feature rankings between both tasks, classification and regression, was very high compared to the other models, with around 55% for all features and 80% for the best representative per cluster (See Supplementary Figure S12). Owing to its capacity to model non-linear relationships, straightforward interpretability, and strong performance in our analyses, we selected random forest as the reference model for assessing feature importance in both classification and regression tasks.

**Table 1.**
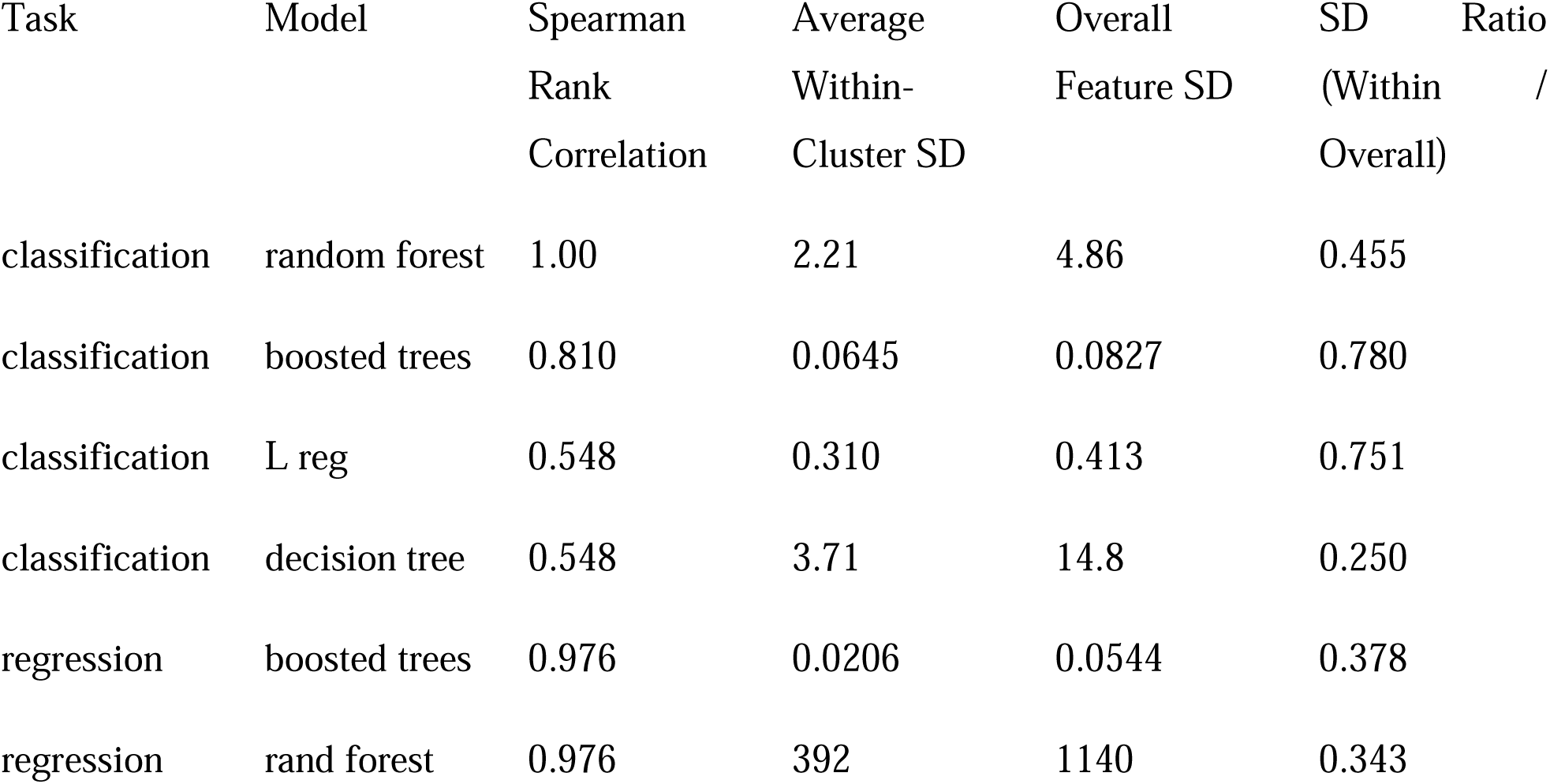
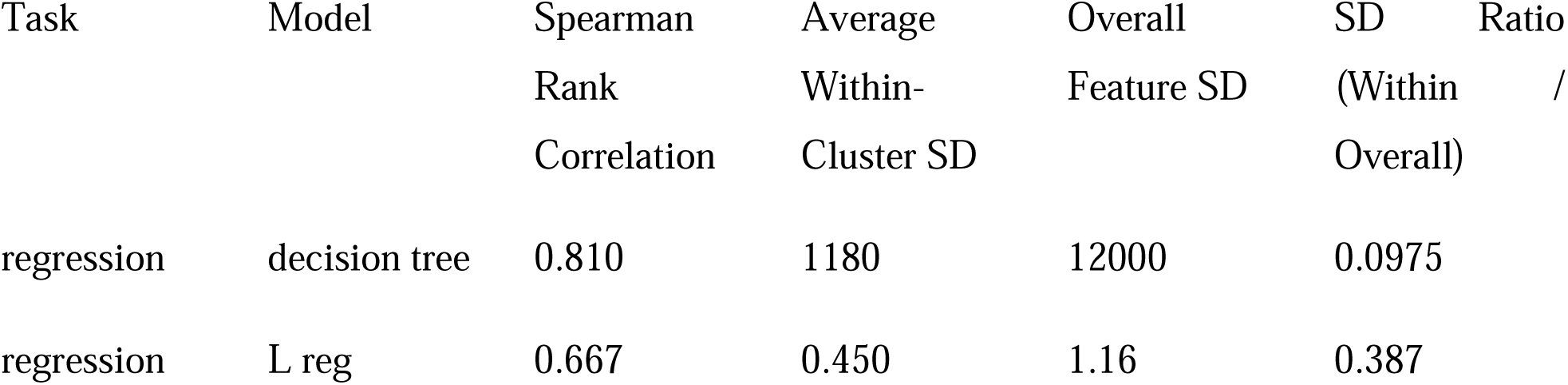
Consistency of feature importance rankings under multicollinearity across four algorithms. Spearman correlation coefficients were calculated using the best representative per clusters between rankings of models developed with all features or with only the best representative per cluster of co-linear features. The table also reports the average standard deviation of feature importance within the same clusters, the average standard deviation across all features, and the ratio between these two values.

### Model performance

Hotspots were predicted with great performance. The RF achieved a mean fold-wise area under receiver operating characteristic curve (AUROC) value of 0.858 (See https://jamonterotena.github.io/bnapus.reco.ml/)—the best among classifiers. The overall AUROC was 0.823 (Supplementary Figure S14), obtained with the probabilities for all original values and their corresponding predictions obtained during cross-validation.

Recombination rate was predicted via regression with moderate performance. RF yielded a mean fold-wise *R*^2^ of 0.662 (See https://jamonterotena.github.io/bnapus.reco.ml/) and an overall *R*^2^ of 0.477, obtained by comparing all the original values with their corresponding predictions (See Figure 5). Nevertheless, predicted versus observed rates were strongly correlated within chromosomes (mean Pearson *ρ* = 0.731), reaching ∼0.90 on C07, C05, A05, and A08 but dropping below 0.50 on C02, A03, C09, and C06 (See Figure 6; See https://jamonterotena.github.io/bnapus.reco.ml/). Thus, although absolute rate estimates were imperfect, the model captured the broad recombination trends along most chromosomes.

**Figure 5.**
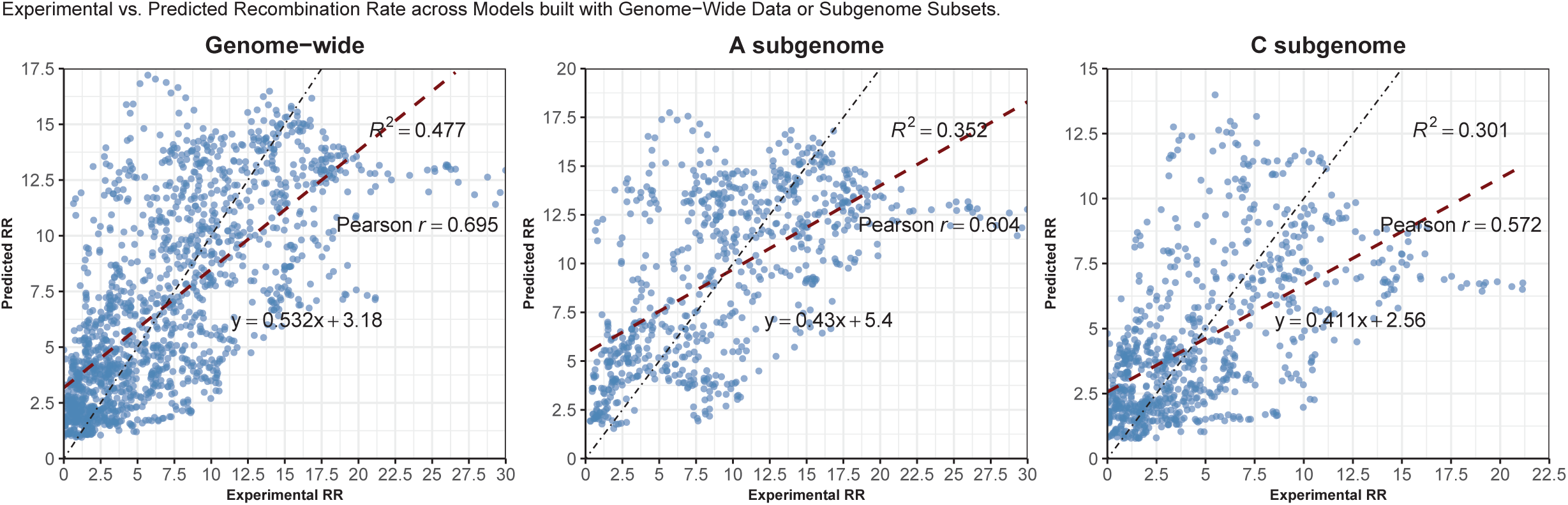
Scatterplot with regression line of experimental and predicted recombination rate values, in cM/Mbp, based on the random forest models built with genome-wide (left), A-subgenome (middle), or C-subgenome (right) data. R2, Pearson p and slope-intercept equations are shown.

**Figure 6.**
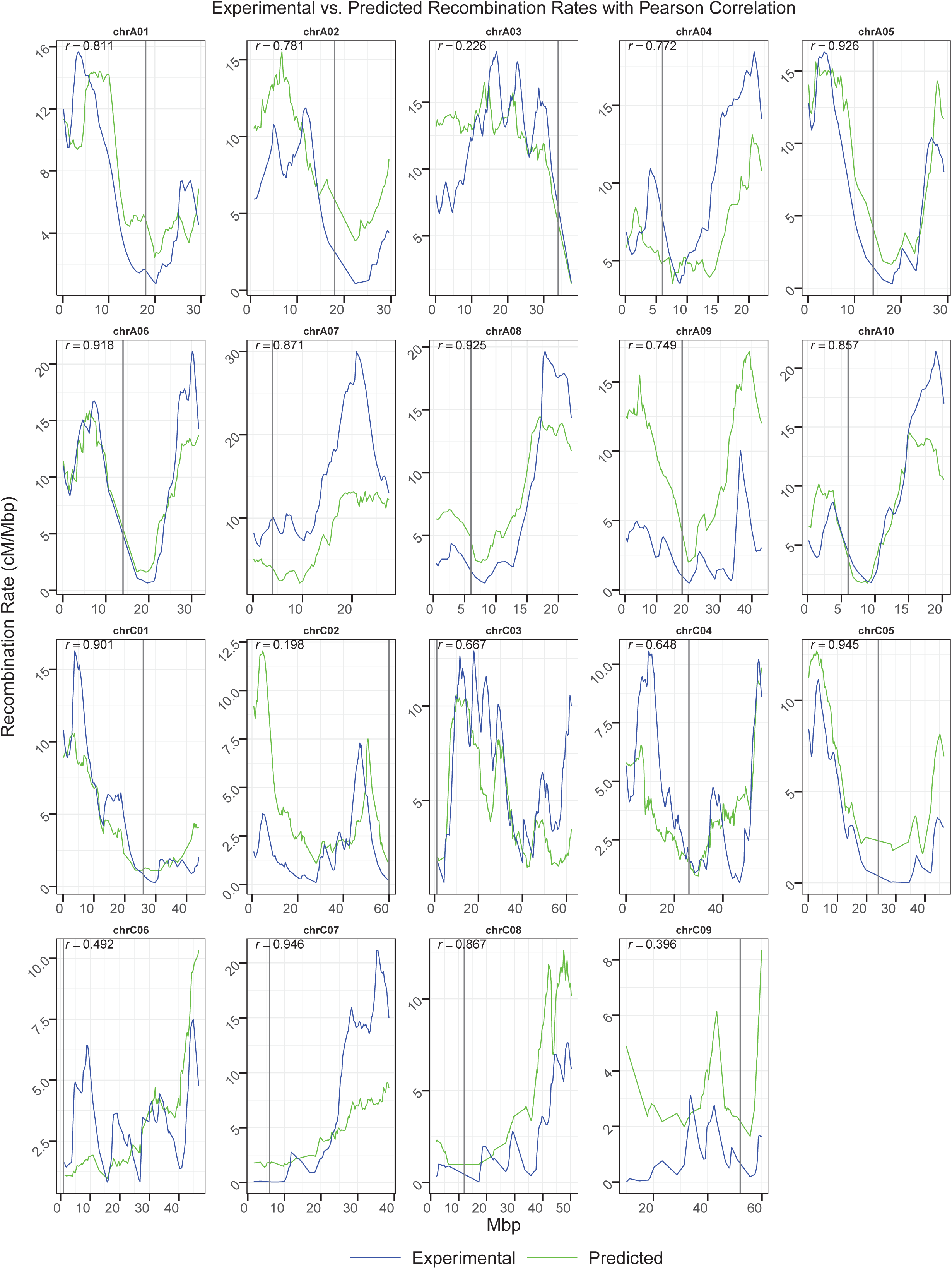
Genome-wide distribution of the actual and predicted values of recombination rate, in cM/Mbp, over 0.3-Mbp genomic bins. Actual and predicted values are indicated by blue and green lines, respectively. Plots are titled with chromosome names. Straight vertical black lines represent centromere locations.

### Feature importance

In the RF regression, CpG methylation was the strongest predictor of crossover rate (See Figure 7). It was followed closely by highly correlated features—gene fraction; CHG, CHH, and TE-body CHH methylation; retrotransposon coverage, gene expression, and transposon coverage. Heterochromatin is enriched for TEs that are transcriptionally silenced by dense CpG/CHG methylation, which is maintained with MET1 or CMT3, respectively. These methyltransferases reinforce marks such as H3K9me2 via RNA-dependent DNA methylation (RdDM), which compact chromatin and suppress COs (Stroud *et al*., 2012; Yelina *et al*., 2015). By contrast, recombination-prone euchromatic arms are TE-poor, transcriptionally active, and hypomethylated, except for local CHH peaks that silence individual TE bodies (Rodgers Melnick *et al*., 2015). Because these features co-localize, their pairwise correlations are intrinsically high, and the RF model distributes importance almost interchangeably across them. Nonetheless, CpG methylation claims the top rank—probably reflecting its dual role as the most direct indicator of TE silencing and the methylation context most tightly linked to crossover suppression. Collectively, these variables provide integrated proxies for local chromatin state, the principal determinant of the recombination landscape (Hsu *et al*., 2022).

**Figure 7.**
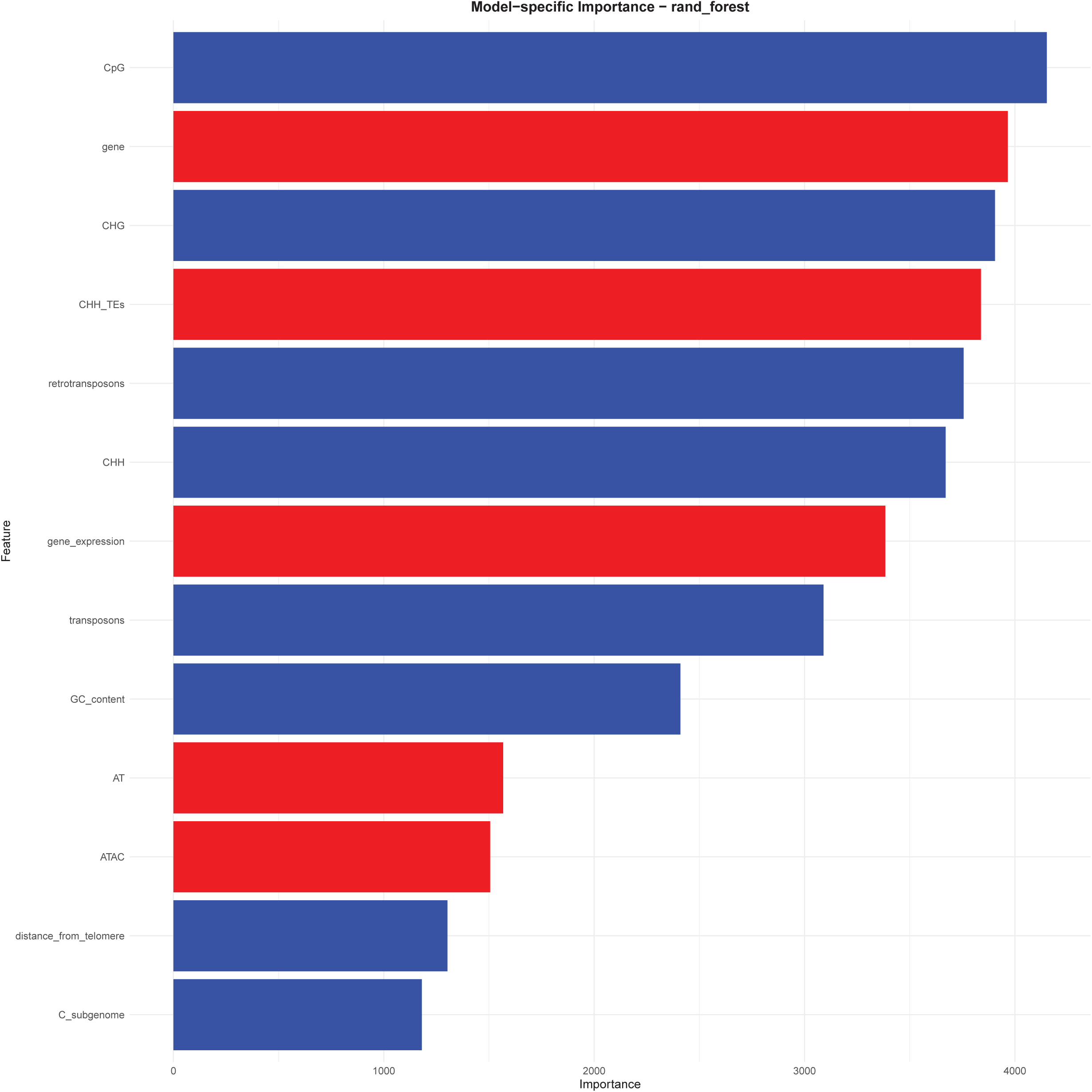
Feature ranking by decreasing Gini impurity-based importance scores obtained by the random forest regression models predicting recombination rates with genome-wide feature data. Features positively correlated with recombination are in red, whereas the negatively correlated ones are in blue.

In classification, feature importance rankings closely resembled those obtained in the regression analysis (See Supplementary Figure S16). Although co-linear features ranked on top, the gap between the leaders and the rest widened. CHG methylation received at least twice the importance score of any feature other than CpG methylation. This suggests that hotspots can be distinguished almost entirely by a narrow set of signals, so adding other, closely related variables yields little additional predictive power.

Telomeric proximity climbed up to the second position in classification even though it ranks among the weakest predictors of absolute recombination rate. Its prominence reflects how the data were sampled: hotspots fall mainly in subtelomeric euchromatin, whereas non-hotspots were drawn largely from pericentromeric heterochromatin (See Figure 2). As a continuous predictor of CO rate, however, telomeric distance loses strength because centromere coordinates are approximate, SNP density is uneven, bins lacking markers were excluded, and the two subgenomes are equally represented, as the C subgenome could have a different recombination landscape, diluting any genome-wide correlation with recombination.

Retrotransposons obtained notably higher importance scores in predicting recombination rate than DNA transposons, even when the two categories are present in comparable numbers. One plausible reason is structural: LTR-Gypsy and LTR-Copia elements—which constitute the bulk of the *Brassica napus* repeatome (Lee *et al*., 2020)—cluster in high number of copies, forming large, dense heterochromatin regions (Wei *et al*., 2013). At the same time, low copy number Copia/Gypsy families are the main CHH-hypermethylated TEs within 1 kb upstream of genes in maize, that coincide with crossover hotspots (Gent *et al*., 2013). Because retrotransposons can mark both large heterochromatic blocks and gene-proximal CHH islands, their genomic coverage may capture recombination-relevant chromatin states more faithfully than the shorter, more scattered DNA-TEs.

### Interaction analysis

After assessing the predictive power of features, we next asked whether their effects are simply additive or if they interacted. Using the RF regression predictions, we assessed how much the interactions between predictors contributes beyond their individual main effects with Friedman’s H-statistic (Friedman and Popescu, 2008). An H value of 0 signifies that interactions between features add no extra predictive information beyond their individual effects, whereas an H of 1 means the model’s signal for those variables is explained entirely by their joint effect, leaving no independent main effects.

Most features showed moderate interaction strengths: on average, the overall interaction of features with every other feature combined explained around 9% of the model’s variance (See Supplementary Figure S18). One pair, however, stood out. Subgenome identity interacted strongly with telomeric distance, accounting for ∼50% of their combined contribution to predicted CO rate, suggesting that contrasting feature trends observed between the A and C subgenome (See Figure 3; See Supplementary Figure S10) significantly shape distinct recombination landscapes between the rapeseed subgenomes.

### Machine learning with single-subgenome input

Given the distinct recombination landscapes of the A and C subgenomes, we asked whether models built and validated solely on one subgenome would outperform those derived from the opposite subgenome or from the genome-wide dataset. Subgenome-specific models may also uncover feature effects that are unique to each subgenomic context.

The RF regression model developed with A subgenome data performed better (mean fold-wise *R*^2^ = 0.597, overall *R*^2^ = 0.352, mean chromosome-wise Pearson *ρ* = 0.780) than its counterpart derived from the C subgenome (mean fold-wise *R*^2^= 0.515, overall *R*^2^= 0.301, mean chromosome-wise Pearson *ρ* = 0.678) (See https://jamonterotena.github.io/bnapus.reco.ml/; See Figure 5). Although the A-specific *R*^2^ is lower than that of the genome-wide model (See Figure 5), its higher chromosome-level correlation shows that it captures spatial trends more faithfully (See https://jamonterotena.github.io/bnapus.reco.ml/). Nearly identical feature-importance profiles in the two subgenomic models indicate that homoeologous chromosomes encode similar recombination patterns, differing mainly in regional signal strength and noise (See Supplementary Figure S19; See Supplementary Figure S20).

ALE curves measure how a single predictor alters the model’s prediction across local intervals while averaging the local values of all other variables, therefore constituting a multicollinearity-robust approach to estimate main effects. Positive ALEs on an interval indicate that the feature raises the predicted recombination rate when moving across the interval, while negative main ALEs represent lowering effects. Overall, ALE profiles correlated well across subgenomes and matched the expected relationship with recombination (See Supplementary Figure S21; See Supplementary Figure S22).

However, ALE profiles for telomeric proximity differed sharply between subgenomes (See Figure 8). For the A subgenome, the most positive local effects appeared in the subtelomeric regions (∼0.25 relative distance from the telomere), where the telomeric proximity increased the predicted recombination rate by up to +0.25. In the C subgenome, the peak shifted inward: pericentromeric segments (0.70–0.80) raised the rate by up to +0.20. Telomeric ends produced the most negative effects in both subgenomes (−0.75 in A, −0.20 in C). Centromeres were strongly suppressive in C but showed a slight positive artefact in A, likely caused by sparse data after excluding SNP-empty bins.

**Figure 8.**
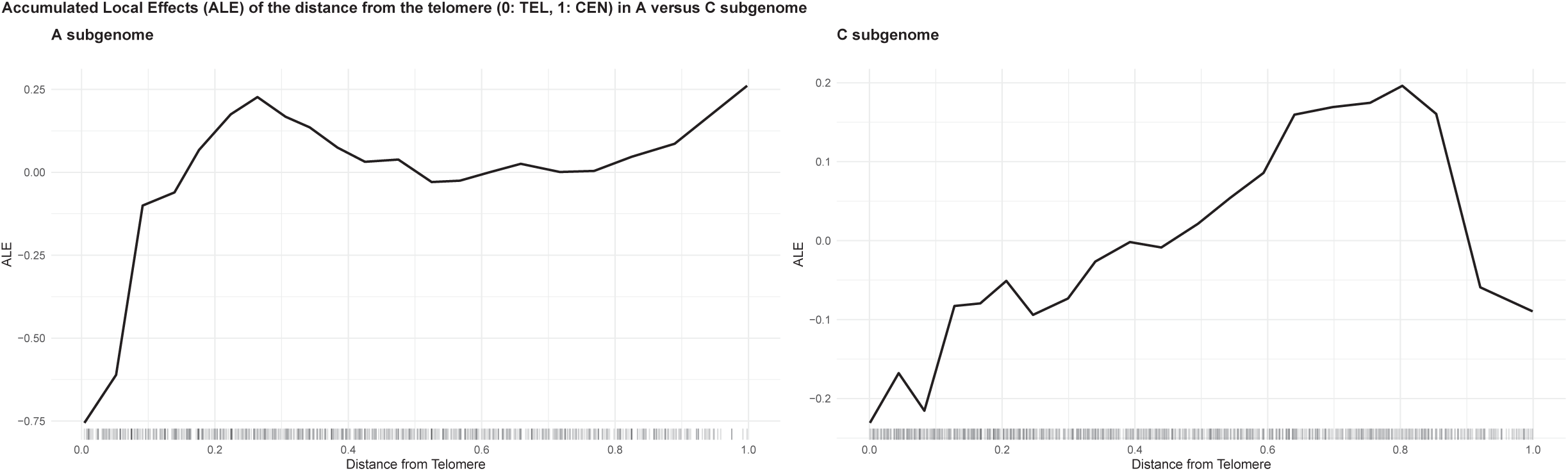
Accumulated local effect (ALE) curves of distance from the telomere to the 0.3-Mbp bins in the models built on A-subgenome data (left) and C-subgenome data (right). The X axis represents the distance from the telomere, ranging from 0 (telomere) to 1 (centromere), whereby the Y axis indicates the ALE, centered around 0. The distributions of bins across the feature values are shown with rugs alongside the X axis.

Taken together, our results show that the two parental subgenomes of rapesed carry distinct recombination landscapes. In the A subgenome (*Brassica rapa*), crossovers cluster toward subtelomeric regions; in the C subgenome (*Brassica oleracea*), they concentrate in pericentromeres. This pattern mirrors earlier genetic maps (Yan *et al*., 2023), where chromosomes C06, C08, and C09 exhibit elevated pericentromeric recombination, whereas their A-homoeologs peak near the telomeres despite being physically shorter. Telomeric distance, however, explains only a small fraction of the variance in crossover rate; the dominant predictors in both subgenomes are the same chromatin-state markers that govern recombination genome-wide.

## Discussion

We built machine-learning models that accurately predicted recombination rate and hotspot location and classified multi-omic features according to their predictive power. We conducted unconventional machine-learning techniques, such as the application of accumulated local effect (ALE) curves to understand the effect that local values had on the target both for single features or combination of feature values. This complements the general effects represented by importance scores, representing a model-agnostic approach to understand model predictions that is strongly robust to multicollinearity. Because co-linear variables are commonplace in biological studies and complicate the estimation of their actual contribution to explaining biological phenomena, we implemented a statistical comparison of results in the presence or absence of multicollinear features across the four algorithms to select the model most consistent in the face of highly correlated features. The random forest was the most consistent model under multicollinearity, reflecting the enhanced robustness of tree-based models compared to linear/logistic regression models (See Supplementary Figure S13). We encourage researchers to incorporate this test when selecting algorithms to ensure fair comparisons among predictors.

The main limitation was the coarse resolution of crossover intervals (See Supplementary Figure S1): low founder diversity and the application of the 15K SNP chip required clustering events into contiguous 0.3-Mbp windows. Whereas most studies use high-density WBS genotyping or 60K SNP arrays in small sets of recombinant inbred line (RIL) to pinpoint breakpoints precisely, we analysed two outbred populations of ∼1,000 plants, detecting far more crossovers and thus bringing recombination-rate estimates closer to reality—after stringent haploMAGIC filtering to remove false positives. Because crossover calls scale with marker density, recombination rates can be confounded by local SNP content; however, after normalising for SNP density, rates remained highly correlated with the raw values, reflecting evidence that hotspots coincide with regions of elevated polymorphism. To avoid such confounding, we excluded SNP density from the predictor set.

The features that indicate chromatin state, well-known to shape recombination globally, are also the major players in *Brassica napus*, suggesting conservation of the main recombination mechanisms. These factors are highly correlated among each other and their tight link with recombination was clear: Proxies for chromatin state modulated CO formation throughout recombination maps (See Figure 1), and consistently obtained highly significant p-values in the t-test (See Supplementary Figure S5), and the highest importance scores by the random forest in predicting recombination rates (See Figure 7) and hotspot locations (See Supplementary Figure S16).

Our results indicate that subgenomes have distinct recombination landscapes in *Brassica napus* (See Figure 8). The A subgenome is subtelomeric-dominant, whereas the C subgenome is pericentromeric-dominant. Besides, the C subgenome was importantly less recombination-prone. These different recombination patterns concord with contrasting feature patterns: The feature profile of the C subgenome—higher for anti-recombinatory factors and lower for the pro-recombinatory ones—evolves differently along chromosome arms, which reflect their distinct recombination landscape (See Figure 3; See Supplementary Figure S10). Changes in the epigenomic landscape after interspecific hybridization between *Brassica rapa* and *Brassica oleracea*, as evidenced by the reported decrease in chromatin-accessible regions in the C subgenome (Quan *et al*., 2025), could have shaped the recombination landscapes differently across subgenomes. Lower recombination frequency could have led to greater conservation of LD in the C subgenome—lower genetic diversity—, intensified with strong selection that is typical of the breeding history of rapeseed (Qian *et al*., 2014).

Our data also show that telomeres display lower recombination, while centromeres lack it altogether (See Figure 1; See Figure 8). Although recombination is generally thought to be high in telomeres, this pattern matches observations in maize, where crossovers peak in subtelomeric regions and decay before the chromosome ends (Rodgers Melnick *et al*., 2015). Because reduced DNA exchange makes these regions less prone to selective sweeps, they may carry a higher genetic load. We therefore recommend breeding strategies aimed at unlocking the latent recombinogenic potential of telomeres in *Brassica napus*.

We encountered contrasting patters between overall CHH methylation rate per bin—negative alongside the CpG and CHG contexts - and methylation of CHHs located in the body of transposable elements - positive. The first finding disagrees with the reported positive associations (Peñuela *et al*., 2022, 2024; Rodgers Melnick *et al*., 2015). However, narrowing CHH methylation to TE bodies results in positive association, possibly reflecting high CHH methylation of TEs in recombination hotspots.

We observed that the coverage with ATAC-accessible regions was positively related with CO formation (See Figure 7). However, we expected this feature to achieve better predictive scores since other features that are universally clearly linked with chromatin accessibility were consistenly among the best predictors. This could be due to the dilution of the ATAC-seq signal. We could not observe clear relationships with nucleotide-based features at this resolution of the recombination data. Therefore, a couple of questions remain inconclusive: Whether hotspots have higher GC content due to GC-biased gene conversions (Rodgers Melnick *et al*., 2015), or if they are enriched for A/T-rich sequences characteristic of gene promoters (Wijnker *et al*., 2013).

## Conclusions

Taken together, our findings show that the principal determinants of crossovers are conserved in *Brassica napus*, explore patterns of multi-omics features previously associated with recombination, and propose a model in which the recombination landscape differs in the two sub-genomes. The subgenome specific patterns could have an impact on allele recombination during breeding and the future efforts to engineer increased recombination rates or altered recombination patterns.

## Materials and methods

### Sequencing data

*Brassica napus* plants belonging to two large multiparental populations were genotyped using a 15K SNP array (Clarke *et al*., 2016). These populations, consisting of 1,573 and 1,550 individuals for population 1 and 2 respectively, originated from two panels of 50 elite German varieties and were developed through four generations of outcrossing following a simulation-based crossing program aimed towards maximizing general combining ability while preserving genetic diversity as described by Krenzer *et al*. (2024). After processing raw SNP data outlined in the ‘Materials and Methods’ section of Abdollahi et al. (Abdollahi *et al*., 2024)), a final dataset of 11,443 SNP markers was retained across all 19 chromosomes of *Brassica napus* (AACC, 2n = 4x = 38). 2,566 individuals were retained—1,243 and 1,323 in population 1 and 2-after discarding plants with missing pedigree information about the founder lines.

### Generation of recombination data

We implemented haploMAGIC (Montero-Tena *et al*., 2024) for detecting recombination events across the two parental haplotypes of the individuals spanning generations G2 to G4. In these generations, the offspring inherited their haplotypes from recombination-informative meioses ocurring in heterozygous parents. haploMAGIC reconstructs parental haplotype sequences using pedigree information by resolving diploid SNP genotypes into phased grandparental haplotype blocks. These blocks become fixed along chromosomes after meiotic recombination, enabling the identification of crossovers within intervals where transitions between parental haplotypes occur. To address generation-specific genotyping error patterns, we configured haploMAGIC with the options min = 2/5/3 for generations G2, G3 and G4, respectively, along with imp = imputeTHonly and cor = correctFalseHom. These settings ensured consistent genome-wide recombination rate trends across generations (See Supplementary Figure S9). A total of 171,276 CO intervals were identified across all informative meioses (median length = 874,808 bp, mean length = 2,626,983 bp) (See Supplementary Figure S1). The link to the raw crossover set can be found in the ‘Data avalability’ section.

We performed two quality filters on the crossover set by (1) removing COs from outlier individuals having more than 100 COs over all meiotic events (See Supplementary Figure S3), and (2) discarding CO intervals longer than the 2 Mbp. The filtered CO set consisted of 148,600 COs(median length = 444,895 bp, mean length = 612,933 bp) (See Supplementary Figure S2).

### Generation of epigenetic feature data

For the epigenetic and transcriptomic features, ATAC-seq and RNA-seq data was generated for 3.5-week-old leaves, 9-day-old roots, immature siliques, 3-day-old seedlings and immature flower buds of rapeseed, while WGBS data was obtained for the first three tissues. Two replicates were sampled from each tissue.

### Calculation of recombination rate

Recombination rates were first estimated by counting the number of crossover events occurring within non-overlapping 0.3-Mbp bins across the genome. When a crossover interval spanned multiple bins, its contribution was proportionally distributed based on the fraction of the interval overlapping each bin. Recombination rate in centiMorgans per megabase (cM/Mbp) was then calculated for each bin using the following formula:

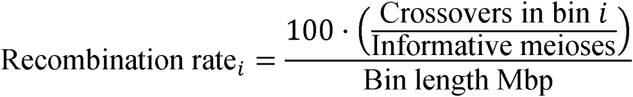

The number of informative meioses varied by population and was equal to twice the number of informative individuals retained after filtering individuals: 1,157 for Population 1 and 1,265 for Population 2.

### Sampling of hotspot and non-hotspot regions

Hotspot bins were defined as those falling within the top 5% of recombination rates across the genome. For each chromosome, we randomly selected the same number of non-hotspot bins (with recombination rate below the 75th percentile, excluding SNP-empty bins) as hotspot bins (above the 95th percentile), thereby ensuring balanced class sizes per chromosome while maximizing the contrast between the two groups (See Figure 2).

### Calculation of features

#### Annotation-based features

Genetic features were incorporated from the *Express 617* genome annotation (Lee *et al*., 2020). Transposable elements (TEs) were annotated using the EDTA pipeline (Su *et al*., 2021), which was provided with a custom TE library. Repeats were subsequently masked using RepeatMasker (Smit *et al*., 2013--2023). Where genomic intervals overlapped between feature types, overlapping regions were trimmed to avoid double-counting.

Transposons were defined as the aggregated fractions of the DNA transposon families DNA/DTA, DNA/DTC, DNA/DTH, DNA/DTM, DNA/DTT, and DNA/Helitrons. Retrotransposons included LTR/Copia, LTR/Gypsy, and LTR/Unknown elements. The overall TE category comprised all annotated transposable element families. For each genomic bin, we calculated the proportion of the genomic bin’s length that was covered by each annotation feature.

#### DNA methylation

Whole-genome bisulfite sequencing (WGBS) data were processed beginning with adapter and quality trimming using TrimGalore 0.6.7 (https://github.com/FelixKrueger/TrimGalore) with the parameters --paired --clip_R1 8 --clip_R2 8 --three_prime_clip_R1 8 --three_prime_clip_R2 8. Trimmed reads were aligned to the *Express 617* v1 reference genome using Bismark 0.24.0. Duplicate reads were removed using the deduplicate_bismark script, and methylation calls were extracted for CG, CHG, and CHH contexts using bismark_methylation_extractor.

Methylation reports were converted into bedGraph format using bismark2bedGraph for downstream processing. Genomic interval coordinates were adjusted to 1-based indexing, and calls from top and bottom strands were merged. For each cytosine position, the single-base methylation rate was calculated as the ratio of methylated calls to total calls. Cytosines called in only a single replicate were excluded. For each tissue, methylation rates were averaged across replicates, and final values were obtained by averaging across all tissues.

TE body CHH methylation (CHH_TEs) represents the single-base methylation rate of cytosines in the CHH context that are located within any annotated transposable element. To obtain the overlap of cytosines and TEs, we applied the intersect function of BEDTOOLS 2.27.1.

#### Dinucleotides

We applied the scripts used in Demirci *et al*. (2018) to calculate the proportion of the bins covered by each dinucleotide.

#### GC content

We used the function nuc of BEDTOOLS 2.27.1 (Quinlan and Hall, 2010) to estimate the proportion of guanine and cytosine bases per bin.

#### Distance from the telomere

We inferred the locations of centromeric regions in the *Express 617* genome using the centromere interval coordinates reported in Mason *et al*. (2016) for the *Darmor* reference genome (Rousseau-Gueutin *et al*., 2020). Centromeric sequences were extracted from the *Darmor* v10 assembly and aligned to the *Express 617* v1 genome using NUCMER 4.0 (Marçais *et al*., 2018) with the options --maxmatch --nosimplify to retain repetitive regions. The resulting alignments were filtered using delta-filter with the options -r -q to retain only the best global hits. Aligned regions consistently clustered in specific genomic areas, allowing us to define a single representative centromeric position per chromosome. In cases where multiple distinct alignment blocks were observed, we prioritized those closest to the centromeric intervals defined by Mason *et al*. (2016).

For each bin, we calculated its relative position along the chromosome arm as the distance from the centromere to the bin midpoint divided by the distance from the centromere to the closest telomere. This ratio ranges from 0 to 1 and reflects how close a bin is to the centromere versus the telomere.

#### Chromatin accessibility

Raw reads were trimmed using cutadapt 4.0 (Martin, 2011) and aligned to the *Express 617* v1 reference genome with Bowtie2 2.4.5 (Langmead *et al*., 2009) using the parameters -k 10 -X 1000 --very-sensitive. Open chromatin regions were identified using Genrich in ATAC-seq mode with the options -j -d 150 -y -a 200 -r.

For each bin, we calculated the overlap with open chromatin regions in each tissue individually. Given the strong tissue specificity of chromatin accessibility, the tissue-level coverage values were first transformed before being averaged across tissues. To constrain the transformed values between 0 (completely inaccessible) and 1 (fully accessible), we applied the following transformation:

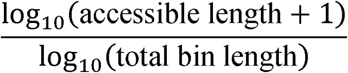

This transformation reduced the skew from large tissue-specific peaks while preserving relative accessibility information across bins.

#### Gene expression

From the RNA-seq data set, trimming was doned using cutadapt 4.0 (Martin, 2011) with the adapter sequence -a CTGTCTCTTATACACATCT. We quantified transcript abundance with kallisto v0.50.0 (Bray *et al*., 2016) using the *Express617* v1 reference. We averaged the transcript per million of genes were averaged across replicates of the same tissue. To account for tissue-specific differences in gene expression, average TPM values within tissues were logarithmically transformed and then averaged across tissues. The formula applied for transformation was:

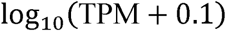

After intersecting these values with the feature table, the gene expression level was calculated as the average TPM among the genes within each bin.

#### Subgenome

A binary feature *C_subgenome*, was created with the values 1 for bins annotated in the subgenome C (chrC01-chrC10) or 0 for bins in the subgenome A (chrA01-chrA09).

#### SNPs

The genomic positions of the SNPs used for crossover detection were used to count the number of SNPs within each bin.

### Construction of feature tables

A feature table was constructed for each machine learning task, with bins represented as rows and the following columns: bin coordinates (chromosome, start, and end base pairs), target value, and feature values. The target variable depended on the task: for classification, the target was a binary variable indicating hotspot presence (TRUE for hotspot bins, FALSE for non-hotspots); for regression, the target was the normalized recombination rate (a continuous numeric value). Feature values were computed per bin using the intersect function of BEDTOOLS 2.27.1 (Quinlan and Hall, 2010), implemented via a custom script (https://github.com/jamonterotena/Custom-BEDTools-intersect). This process extracted all feature elements overlapping with the bin intervals. The link to the feature table can be found in the ‘Data availability’ section.

### Two sample statistical tests

We applied a two-sample Student’s *t*-test with equal variance to compare the means of each feature between hotspot and non-hotspot regions. The two-sample chi-square test for categorical values was applied to test differences between the number of hotspots between the A and C subgenomes. The null hypothesis (H_0_) assumed no significant difference in the mean values of a given feature between the two groups, while the alternative hypothesis (H_1_) proposed that a significant difference existed. In order to meet assumptions of the *t*-test, we normalized them by subtracting the mean and dividing by standard deviation. To account for the increased probability of false positives due to multiple comparisons, we applied the Bonferroni-Hochberg correction to adjust the Ρ-values and control the false discovery rate (FDR). We considered features with adjusted Ρ-values below 0.1 were considered significant.

### Simple exponential smoothing

Before inputting the data for training predictive models, exponential smoothing with *alpha* = 0.1 was applied using the function ses of the R package forecast 8.24.1 (Hyndman and Khandakar, 2008) to the recombination rate values and the feature values to remove noise associated with the abrupt change in adjacent windows. We did not smooth features in which genome splitting could not cause noise, e.g. the distance from the telomere and C subgenome.

### Machine learning modeling

Before modeling, we grouped homoeologous chromosomes together during cross-validation in order to prevent data leakage from shared evolutionary history. Chromosome A10, which lacks a homoeologous counterpart in the *Brassica napus* genome, was grouped with chromosomes A09 and C09 due to their reported chromosomal synteny (Higgins *et al*., 2018). This resulted in a 9-fold cross-validation strategy, where each chromosome group served as the validation set in one fold, while the remaining groups were used for training. Homoeologs were replaced by chromosomes when models were developed using data from a single subgenome. Bins were shuffled within chromosomes to remove positional dependencies and ensure that models relied solely on feature values. During each fold of cross-validation, data preprocessing was applied to the training set, including conversion of categorical variables into dummy variables, removal of predictors with zero variance, and normalization of all numeric predictors. Model parameters were then tuned using Bayesian optimization to improve predictive performance, and models were evaluated on the validation fold using metrics appropriate to the task.

Machine learning models were developed to classify recombination hotspots and to predict normalized recombination rates. Modeling was performed in R using the tidymodels framework (Kuhn and Wickham, 2020). For each task, we trained four algorithms: decision trees (DT), regularized logistic (or linear) regression (LR), random forests (RF), and boosted trees (BT). Hyperparameters were optimized for each model type. DT models included tuning of cost complexity, tree depth, and minimum node size. LR models were tuned for penalty and mixture parameters. RF models were optimized for the number of predictors sampled at each split (mtry), number of trees, and minimum node size. BT models were tuned for mtry, number of trees, learning rate, tree depth, loss reduction, and minimum node size. For classification, the model achieving the highest average ROC AUC across cross-validation folds was selected. For regression, the best model was selected based on the highest average *R*^2^ . Additionally, the Pearson’s correlation coefficient was calculated by chromosomes between predicted and actual values.

### Feature importance estimation

To assess the contribution of individual features toward predicting hotspots or recombination rate, we evaluated model-specific importance metrics. In DT and RF models, importance was quantified as the total reduction in Gini impurity (for classification) or in residual sum of squares (for regression) associated with splits on each feature. In BT models, importance was measured as the total gain, representing the cumulative improvement in the loss function attributed to each feature across all trees. For LR models, feature importance was derived from the absolute value of standardized coefficients, reflecting the magnitude and direction of each predictor’s contribution after penalization.

Feature direction was derived differently for linear and tree-based models. For LR models, direction corresponded to the sign of the standardized coefficient, indicating whether the predictor increased (positive sign) or decreased (negative sign) the likelihood of hotspot occurrence or recombination rate. For tree-based models, direction was inferred post hoc: in classification tasks, it was based on whether the mean feature value was greater in predicted hotspot bins than in predicted non-hotspot bins, while in regression tasks, it was determined by the sign of the Spearman correlation between feature values and the model-predicted recombination rate.

### Model selection

Among all the algorithms tested, we selected a model for further analyses based on two criteria: (1) its predictive performance, determined by the best results obtained during Bayesian hyperparameter optimization, and (2) the robustness of its feature importance rankings under multicollinearity.

We assessed this robustness by comparing models trained on two feature sets: one containing all features, and another containing a single representative per predefined feature cluster. Clusters grouped highly correlated features (Pearson correlation coefficient > 0.80) (See Supplementary Figure S11), including one cluster with general methylation rate features (CpG, CHG, CHH), and annotation-based features (gene and retrotransposon coverage), and another cluster comprising nucleotide-based features such as GC content and AT dinucleotide frequency. In the reduced feature set, we retained only the representative of each group—namely, CpG for methylation/annotation-based features and AT for nucleotide composition—instead of all correlated features.

For each combination of model, feature set, and machine learning task (classification or regression), feature importance scores were computed and used to rank features according to their relevance to the model. To quantify the consistency of feature importance rankings between the two feature sets, we calculated the Spearman rank correlation per model and task. Additionally, we evaluated how each model handled co-linear features by computing the average standard deviation of importance scores within multi-feature clusters, divided by the overall standard deviation across all features. Finally, Spearman correlations between ranking outputs were also computed across tasks and models to assess consistency across configurations.

### Quantification and visualization of feature effects and interactions

To investigate the contribution and interactions of the analyzed features in our predictive regression models of normalized recombination rate, we employed H-statistics to quantify interaction strength between features, and Accumulated Local Effects (ALE) plots to visualize both individual and joint effects in a model-agnostic framework. All computations and visualizations were performed using the iml package in R (Molnar, 2018).

## Author contribution

JAM: performed research, wrote the code, wrote the manuscript. SFZ: performed research, generated data, edited manuscript. GY: assisted in the analysis. TK: performed research. AB: performed research. RJS: provided critical comments, edited manuscript. AAG: conceived research, supervised, edited manuscript, acquired funding.

## Data availability

Data generated in this study has been made available at: https://osf.io/362d8/

The R code used to reproduce the genome-wide, A subgenome, and C subgenome analyses is available at: https://jamonterotena.github.io/bnapus.reco.ml/

## Acknowledgements

Analysis by JAM was supported by grant 490622210 from the German Research Foundation to AG. Data generation and analysis by TK, AA and RJS were supported by grant 031B0187 from the German Federal Ministry of Education and Research (BMBF) within the project BreedPatH. This project was supported by the LOEWE Start Professorship to AG from the Hessian Ministry of Higher Education, Research, Science and the Arts.

## Abbreviations

ALE: accumulated local effects
AT dinucleotide: adenine–thymine dinucleotide
AUROC: area under the receiver operating characteristic curve
bp: base pair
cM/Mbp: centimorgan per megabase pair
CO: crossover
CpG: cytosine–phosphate–guanine (CG)
DNA/DTA: hAT transposons
DNA/DTC: CACTA transposons
DNA/DTH: Helitron transposons
DNA/DTM: Mutator transposons
DNA/DTT: Mariner/Tc1 transposons
DT: decision tree
FDR: false discovery rate
G (in CpG and CHG): guanine
GB: gradient boosting
GC content: guanine–cytosine content
H (in CHH and CHG): any nucleotide except guanine
Kbp: kilobase pair
LR: linear/logistic regression
LTR: long terminal repeat
Mbp: megabase pair
P: P-value
ρ: correlation coefficient
RdDM: RNA-dependent DNA methylation
RF: random forest
RIL: recombinant inbred line
R²: coefficient of determination
ROC: receiver operating characteristic
SD: standard deviation
siRNA: small interfering RNA
SNP: single nucleotide polymorphism
TE: transposable element
TPM: transcripts per million

## Supplementary legends

**Supplementary Figure S1**. Distribution of raw crossover (CO) interval lengths detected with haploMAGIC in two large rapeseed multiparental populations. The x-axis represents the CO interval size in megabase pairs, and the y-axis shows the number of crossovers. The vertical solid red line indicates the median, while the vertical dashed red line denotes the mean.

**Supplementary Figure SS2**. Distribution of crossover (CO) interval lengths detected with haploMAGIC in two large rapeseed multiparental populations after applying filtering criteria. The x-axis represents the CO interval size in megabase pairs, and the y-axis shows the number of crossovers. The vertical solid red line indicates the median, while the vertical dashed red line denotes the mean.

**Supplementary Figure S3**. Distribution of the total number of crossovers per individual. The x-axis indicates the total number of crossovers per individual across paternal and maternal meiosis and all chromosomes, while the y-axis shows the number of individuals. The vertical dashed red line marks the filtering threshold of 100 crossovers.

**Supplementary Figure S4**. Genome-wide recombination rate (in cM/Mbp) by population over 0.3-Mbp genomic bins, with Pearson correlation coefficients reported for each chromosome. Populations are represented by different colors. Chromosomes are arranged column-wise by subgenome and row-wise by chromosome number. Centromere locations are indicated with thick dark grey vertical lines.

**Supplementary Figure S5**. Feature significance ranking based on log_10_ P-values from two-sample statistical tests. The C subgenome values were obtained using the chi-square test, whereas all other values were derived from the t-test. The vertical dashed line marks the significance threshold (P = 0.1). Red bars represent features enriched in hotspots, whereas blue bars represent features depleted in hotspots.

**Supplementary Figure S6**. Genome-wide recombination rate (in cM/Mbp) per SNP marker and feature trends over 2-Mbp genomic bins. The grey areas represent recombination rate normalized by SNP marker number, with the overlaid colored line showing its value. Features include CpG methylation rate, transposable element (TE) fraction, gene fraction, gene expression level (log_10_ TPM + 0.1), and SNP content, each normalized by their genome-wide maximum and represented by light blue, purple, red, pink, and green lines, respectively. Chromosomes are arranged column-wise by subgenome and row-wise by chromosome number. Centromere locations are indicated with thick dark grey vertical lines.

**Supplementary Figure S7**. Chromosome-wise mean number of crossovers detected by population. The x-axis represents generation, the y-axis shows the mean total number of crossovers, and colors indicate different populations.

**Supplementary Figure S8**. Chromosome-wise SNP density in number of SNPs per megabase pair (Mbp). The x-axis shows chromosome names, and the y-axis shows SNP density per Mbp. Light green bars represent the A subgenome, and dark green bars represent the C subgenome. Density values are indicated above each bar. Additional information, including total SNP counts and mean chromosome size per subgenome, is provided in the upper right corner of the figure.

**Supplementary Figure S9**. Number of crossovers detected with haploMAGIC under the min2/5/3 parameter setting, separated by generation.

**Supplementary Figure S10**. Density (left) and distribution along chromosome arms (right) of feature values in the A (orange) and C (blue) subgenomes for retrotransposon coverage; CpG, CHG, CHH, and TE body CHH methylation; gene expression; gene coverage; and recombination rate. In the scatterplots, the x-axis represents distance from the telomere, and the y-axis shows feature values. LOESS regression lines are fitted for each subgenome. Recombination rate values above 1 cM/Mbp have been removed for clarity.

**Supplementary Figure S11**. Pairwise relationships among genomic features. The lower triangle displays scatterplots, the upper triangle reports Pearson correlation coefficients (r) with significance (*** P < 0.001, * P < 0.01, * P < 0.05), and the diagonal shows the univariate distributions of each feature. A and C subgenomes, labelled in the diagonal cell corresponding to subgenome, correspond to values 0 and 1 respectively.

**Supplementary Figure S12**. Spearman rank correlation in feature importance rankings between classification and regression tasks for four algorithms when all features were present (blue) and when only the best representative per cluster was included (red).

**Supplementary Figure S13**. Heatmap of Spearman correlation coefficients in feature importance rankings across the four machine learning algorithms used in the analysis.

**Supplementary Figure S14**. Receiver operating characteristic (ROC) curve of the random forest classifier developed using genome-wide data for hotspot prediction. The curve was constructed using the original feature values and estimated probabilities. The area under the ROC curve (AUROC) is annotated.

**Supplementary Figure S15**. Feature rankings by decreasing importance scores obtained from each regression model (top left: boosted trees; top right: decision tree; bottom left: linear regression; bottom right: random forest) predicting recombination rates with genome-wide data. For the random forest, boosted trees and decision tree models, features positively correlated with predicted recombination rates are shown in red, while negatively correlated features are shown in blue. For the logistic regression model, colors correspond to the sign of the coefficients of terms assigned by the model.

**Supplementary Figure S16**. Feature ranking by decreasing Gini impurity-based importance scores obtained from the random forest classifier in the hotspot prediction task using genome-wide data. Features that are more abundant in the hotspots shown in red, while those more abundant in the non-hotspots are shown in blue.

**Supplementary Figure S17**. Feature rankings by decreasing importance scores obtained from each classifier (top left: random forest; top right: boosted trees; bottom left: logistic regression; bottom right: decision tree) predicting hostpot location with genome-wide data. For the random forest, boosted trees and decision tree models, features that are more abundant in the hotspots are shown in red, while those more abundant in the non-hotspots are shown in blue. For the logistic regression model, colors correspond to the sign of the coefficients of terms assigned by the model. Feature ranking by importance scores obtained from each algorithm in the classification task of predicting hotspots with genome-wide data. Features positively correlated with recombination are shown in red, while negatively correlated features are shown in blue.

**Supplementary Figure S18**. Features ranked by decreasing interaction strength, as measured by Friedman’s H-statistic.

**Supplementary Figure S19**. Feature ranking by decreasing Gini impurity-based importance scores obtained by the random forest regression models predicting recombination rates with A-subgenome feature data. Features positively correlated with recombination are in red, whereas the negatively correlated ones are in blue.

**Supplementary Figure S20**. Feature ranking by decreasing Gini impurity-based importance scores obtained by the random forest regression models predicting recombination rates with C-subgenome feature data. Features positively correlated with recombination are in red, whereas the negatively correlated ones are in blue.

**Supplementary Figure S21**. Accumulated local effect (ALE) curves for features in the model built using A-subgenome data. The x-axis shows feature values, and the y-axis shows the ALE, centered around zero.

**Supplementary Figure S22**. Accumulated local effect (ALE) curves for features in the model built using C-subgenome data. The x-axis shows feature values, and the y-axis shows the ALE, centered around zero.

