## Supplementary Figure for "Machine learning and multi-omic analysis reveal contrasting recombination landscape of A and C subgenomes of winter oilseed rape"

Supplementary Figure 1

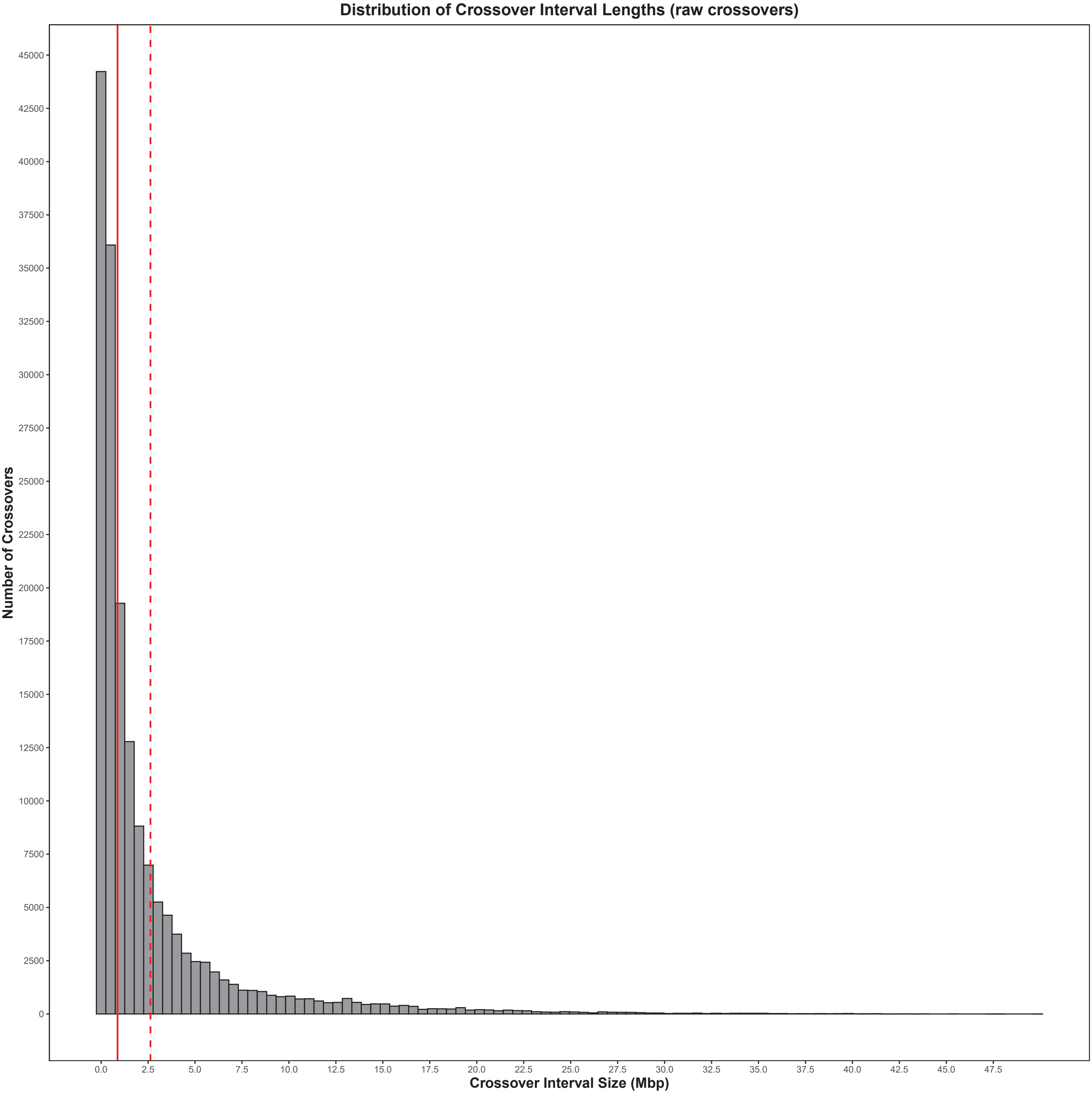

### Supplementary Figure 2

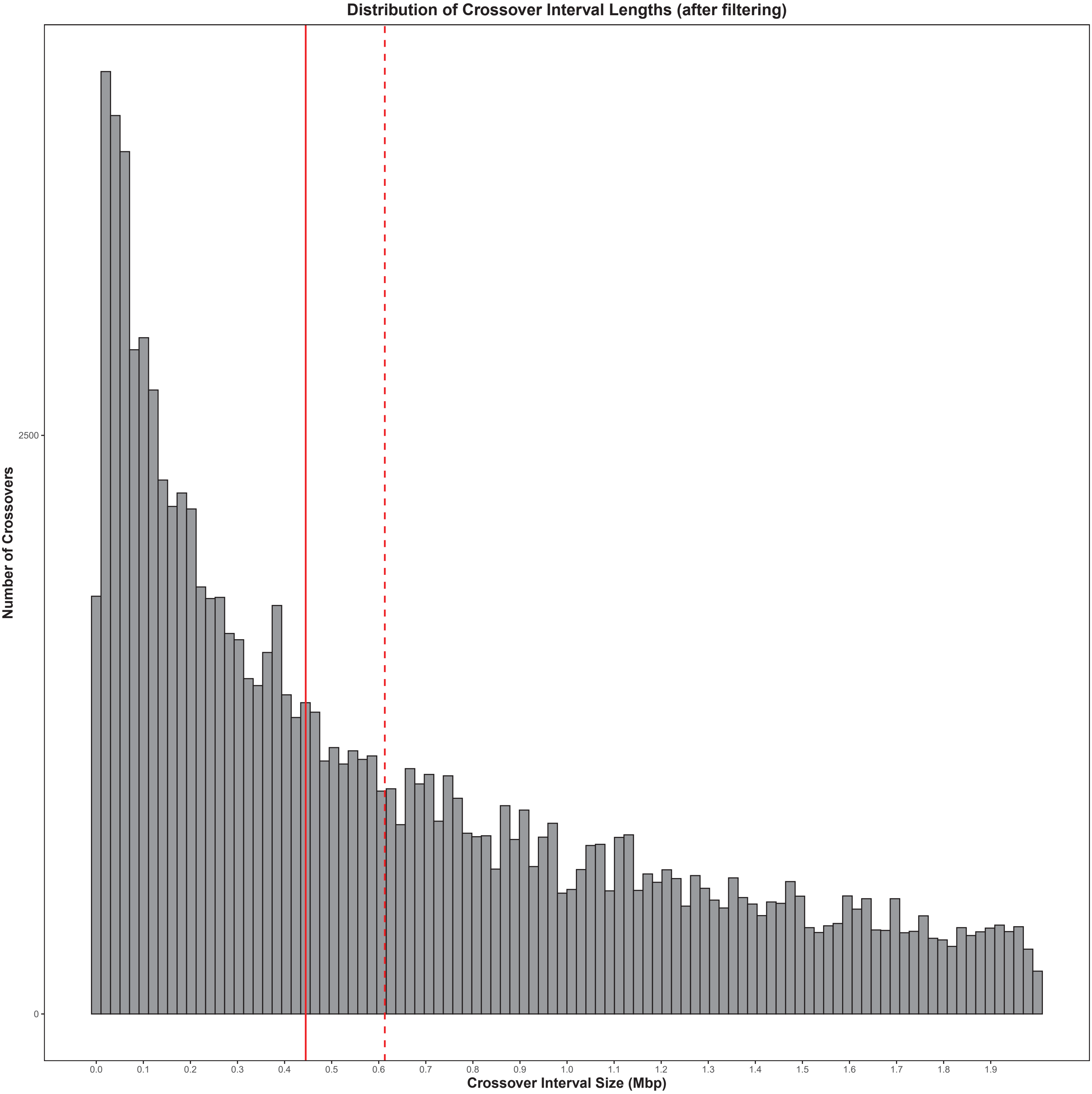

Supplementary Figure 3

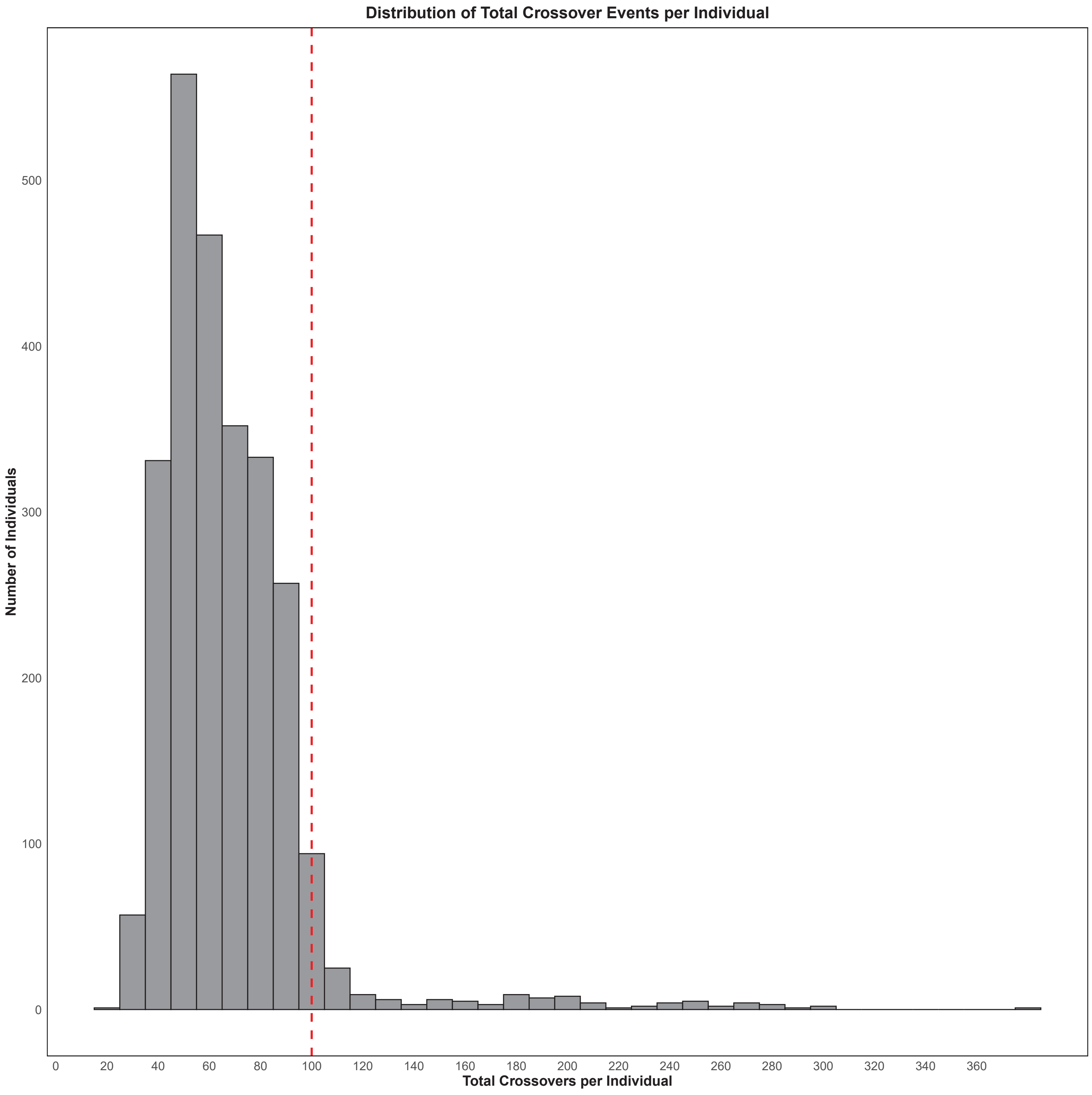

Supplementary Figure 4

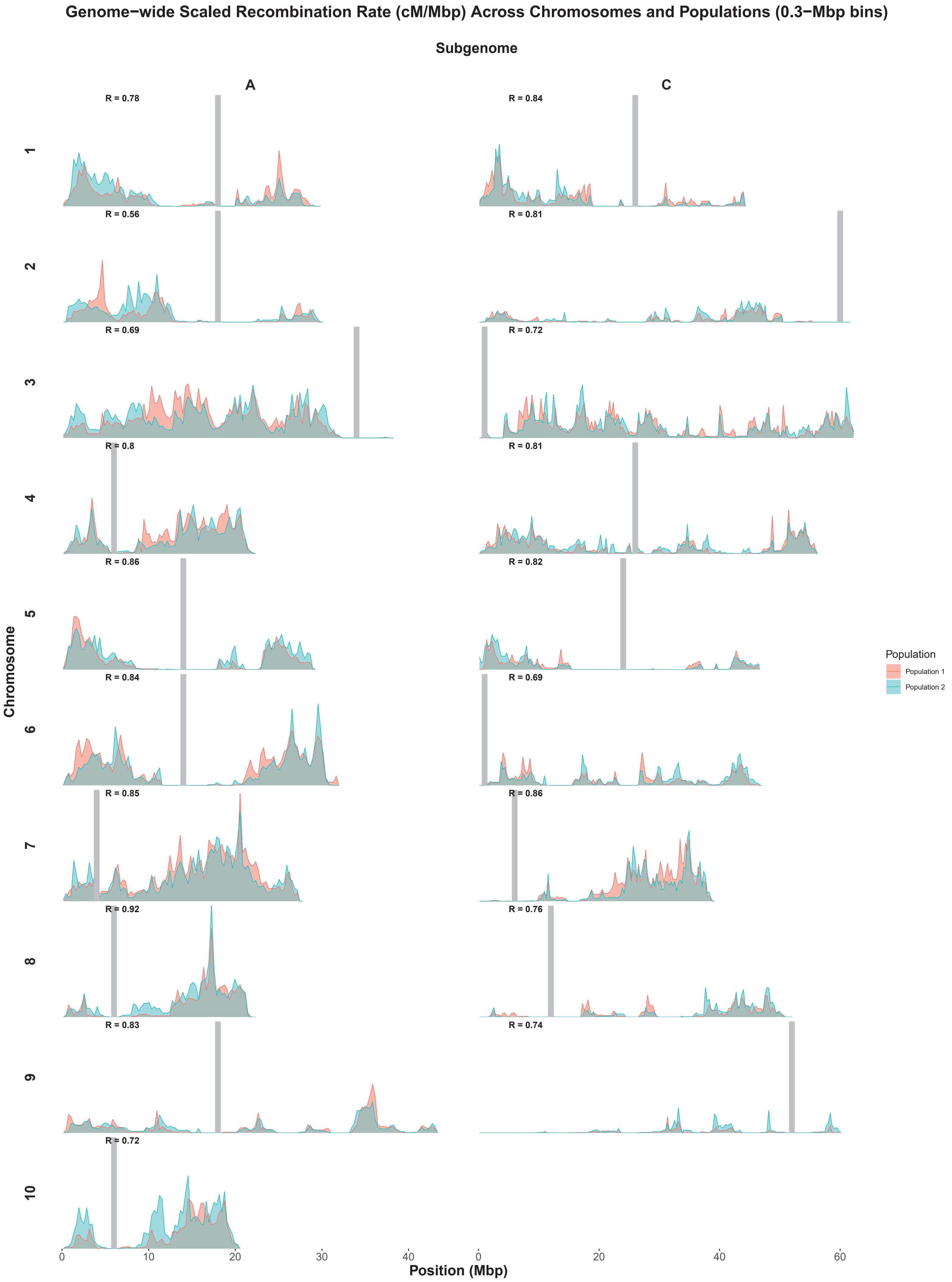

### Supplementary Figure 5

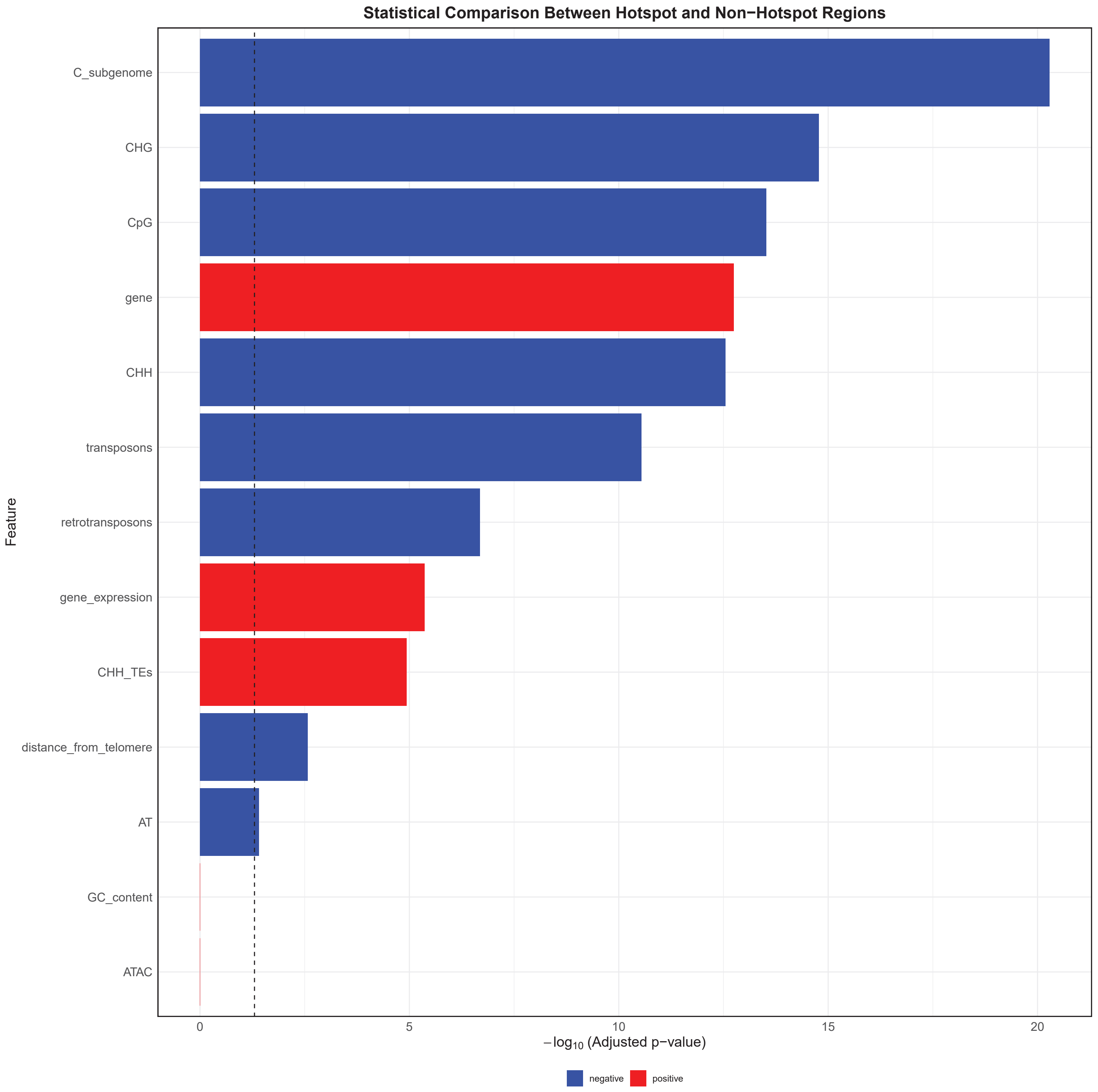

Supplementary Figure 6

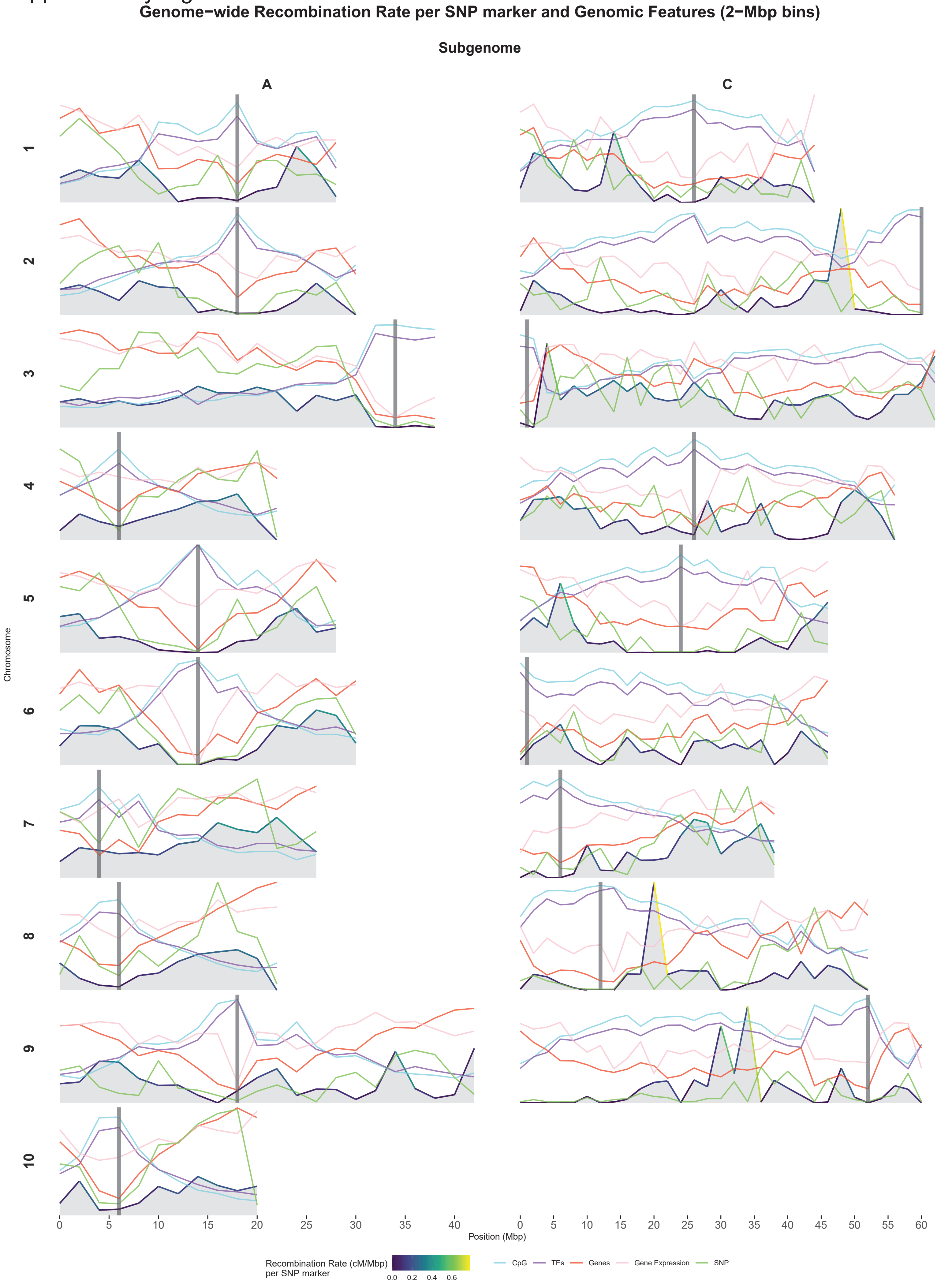

### Supplementary Figure 7

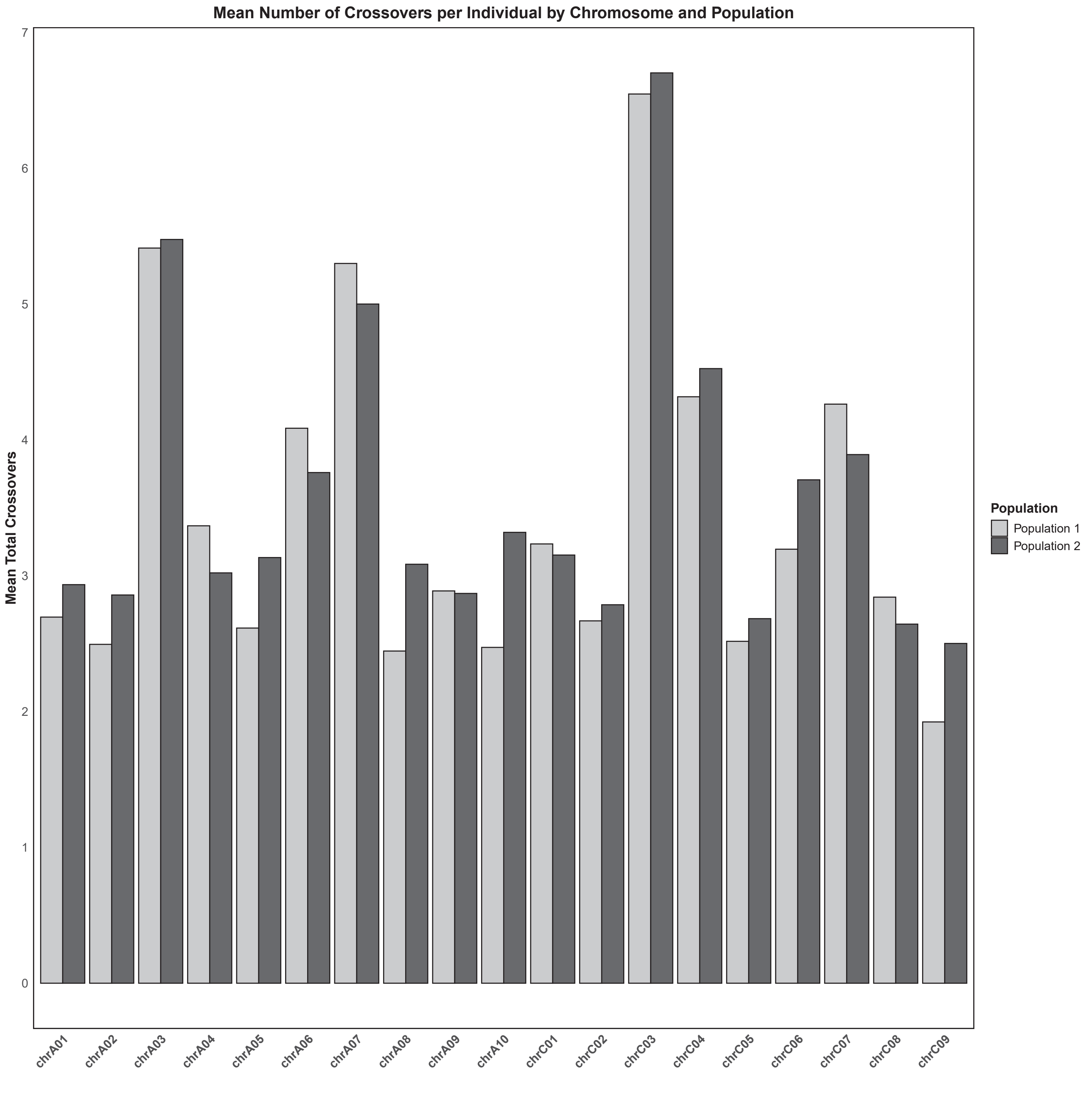

### Supplementary Figure 8

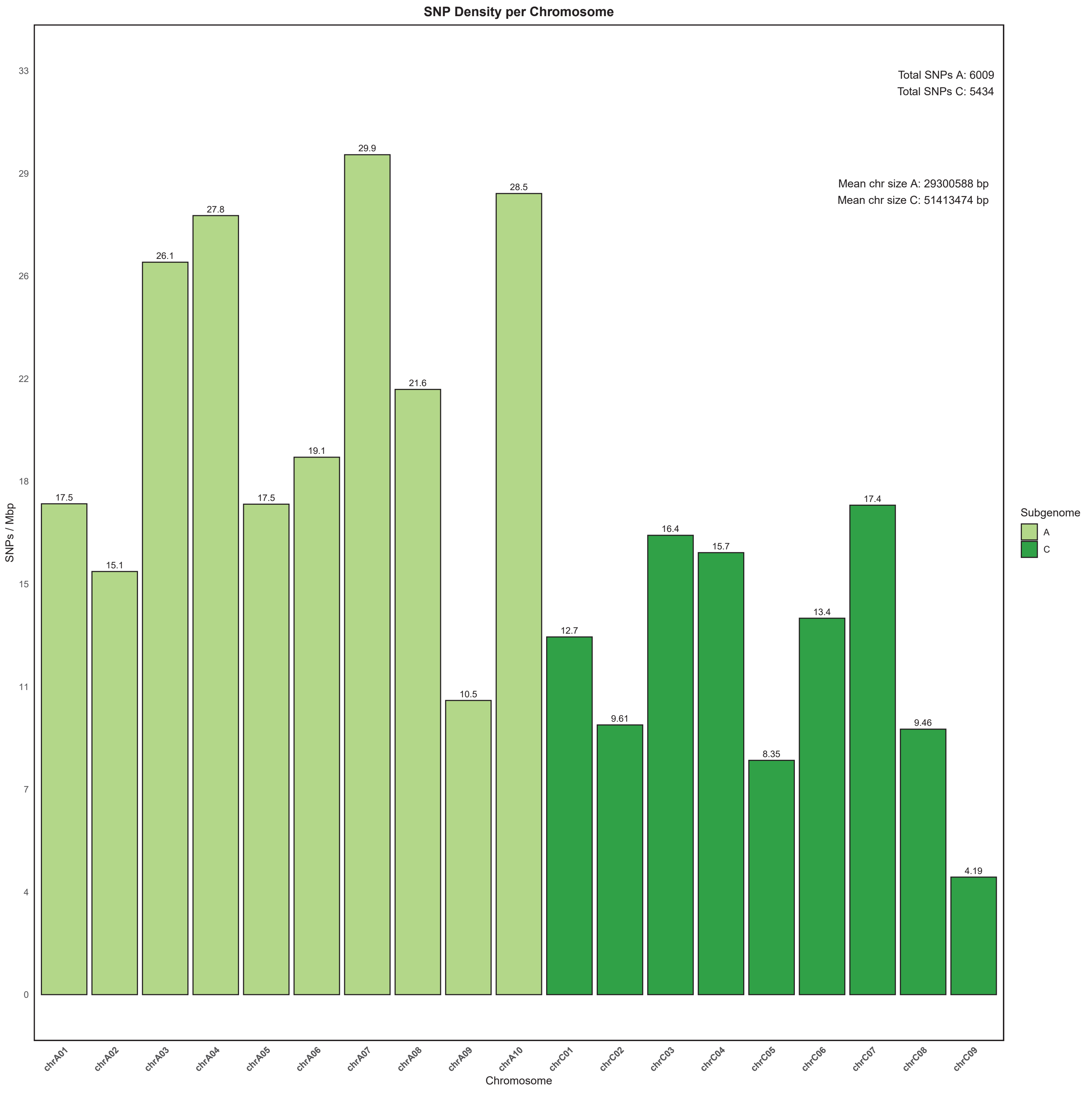

Supplementary Figure 9

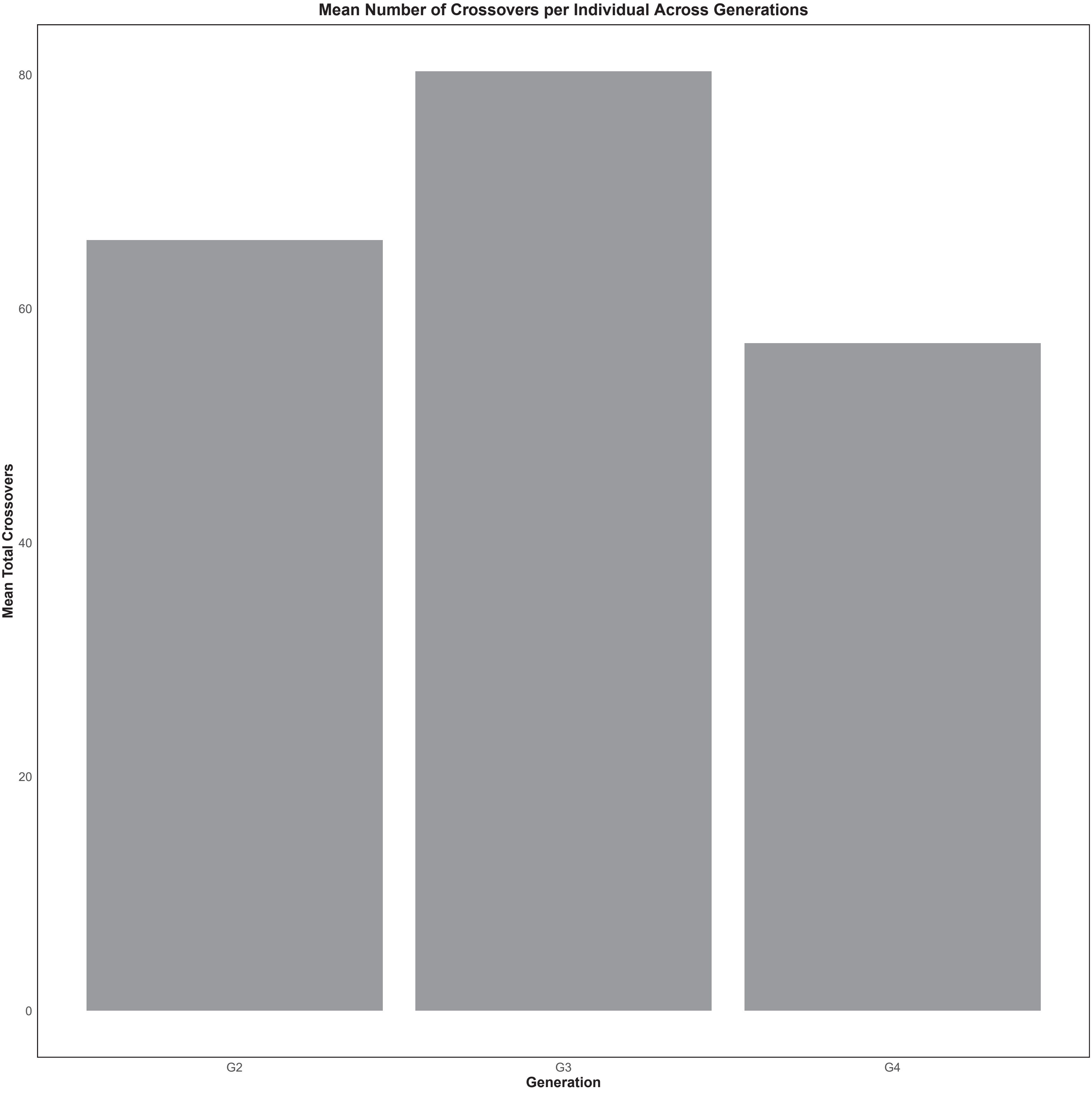

### Supplementary Figure 10

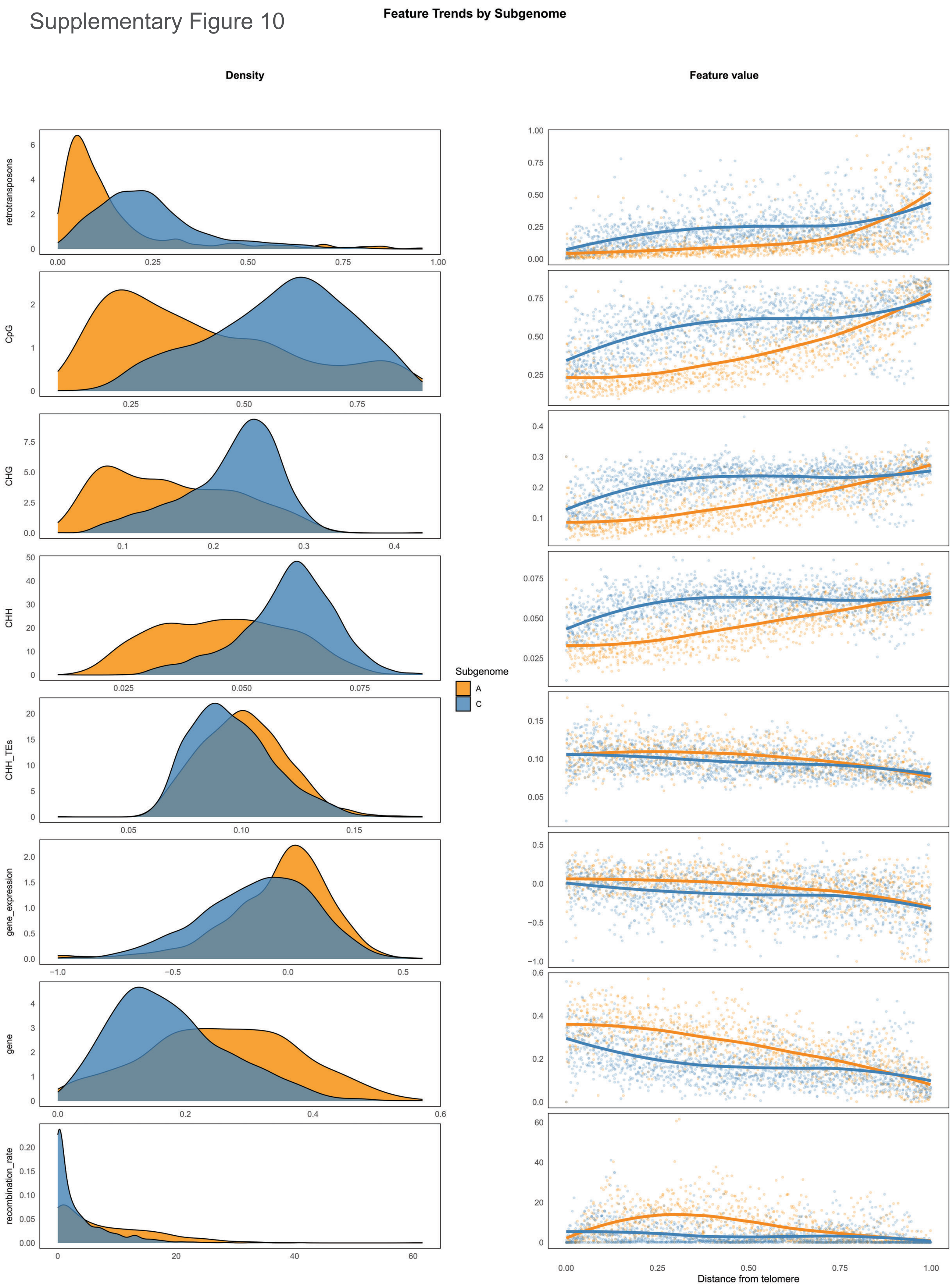

### Supplementary Figure 11

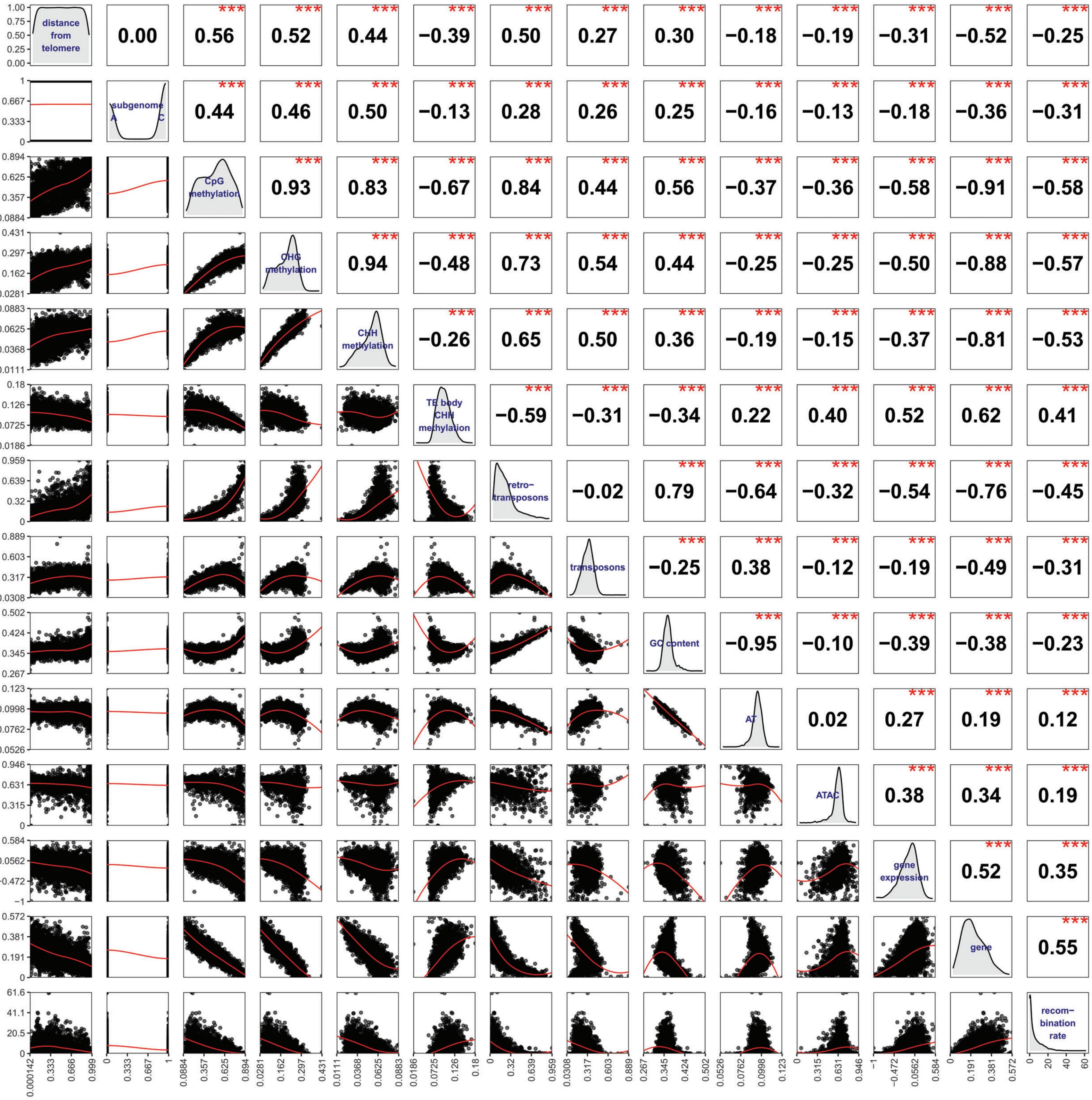

### Supplementary Figure 12

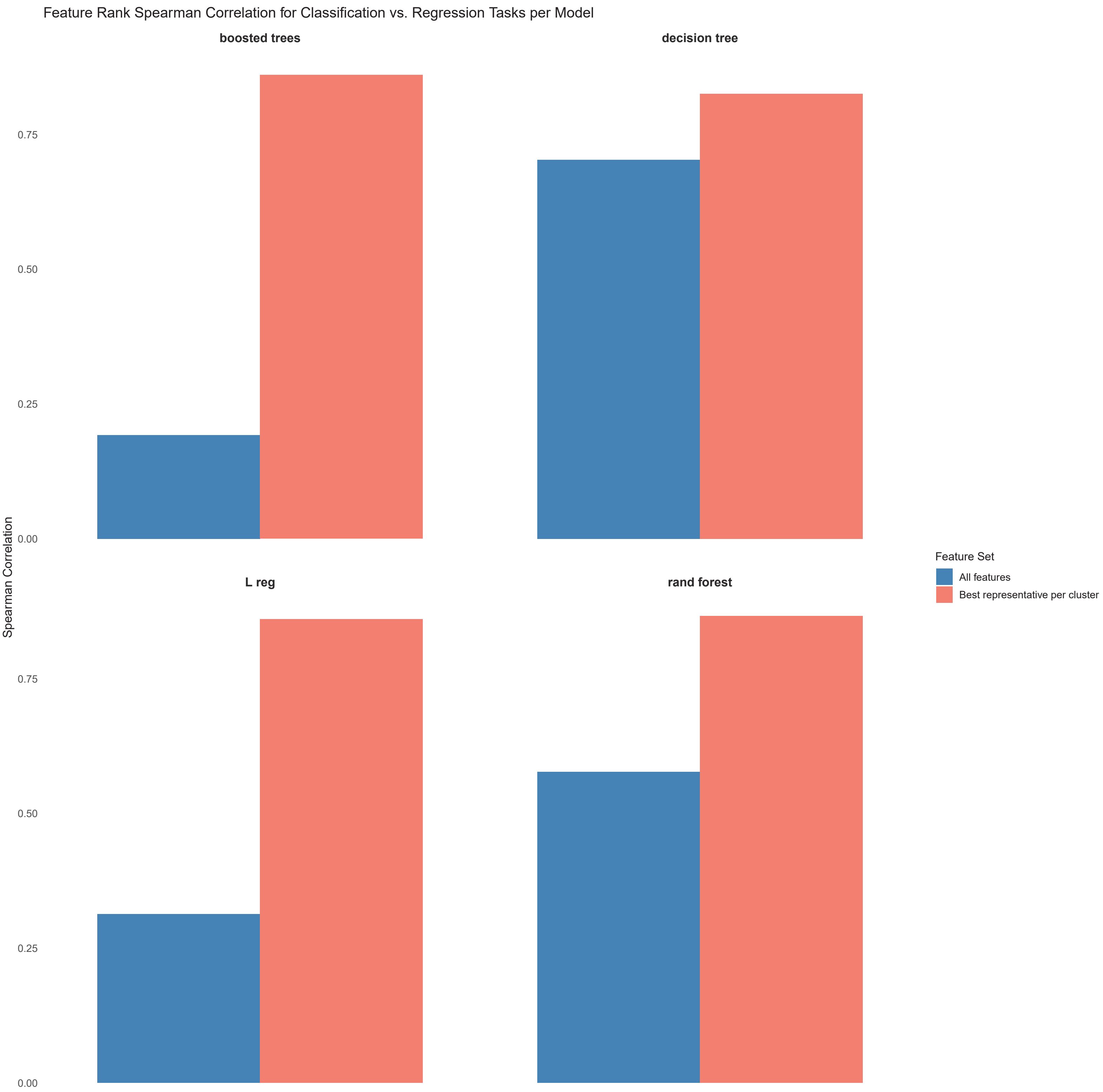

### Supplementary Figure 13

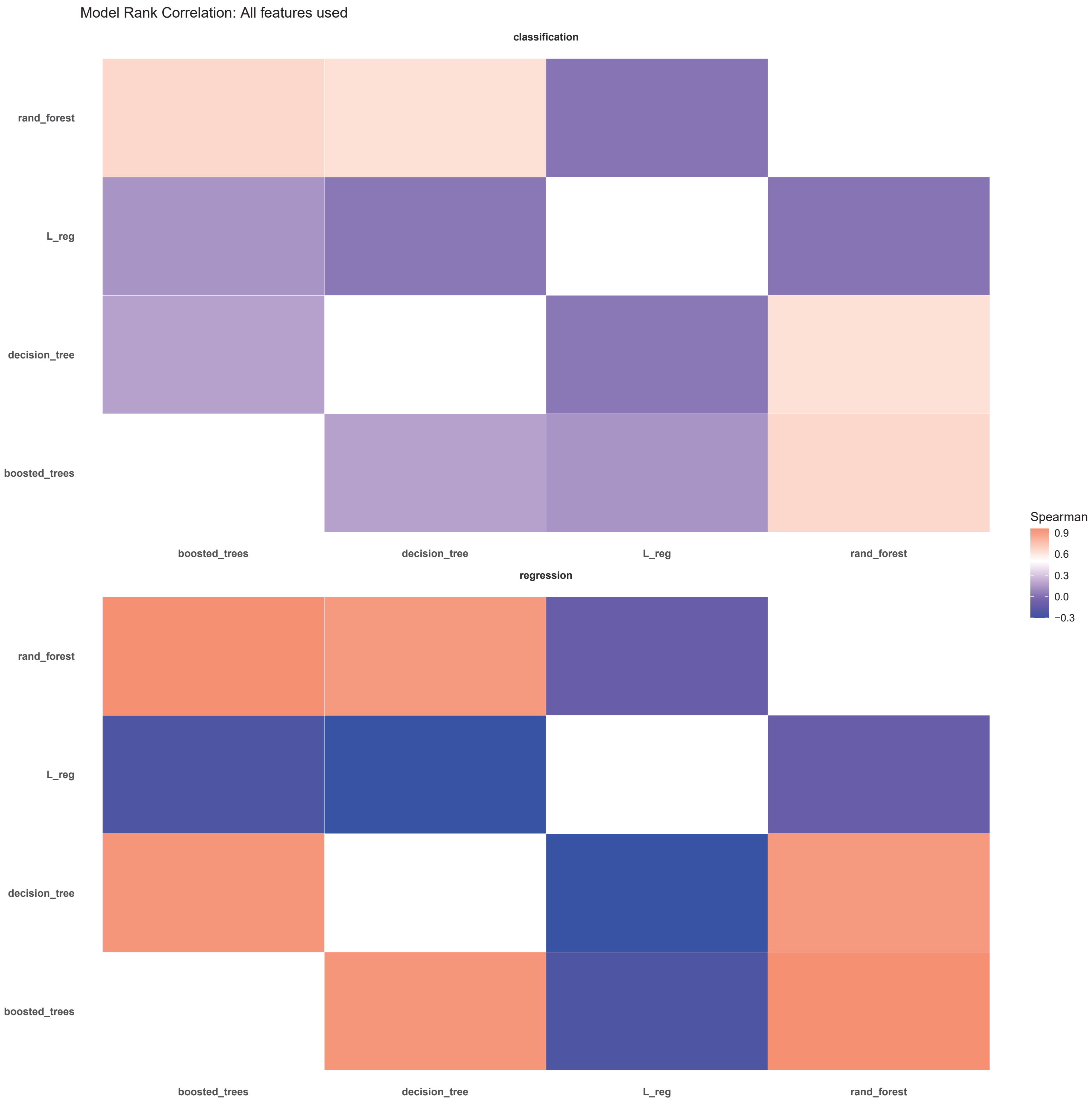

Supplementary Figure 14

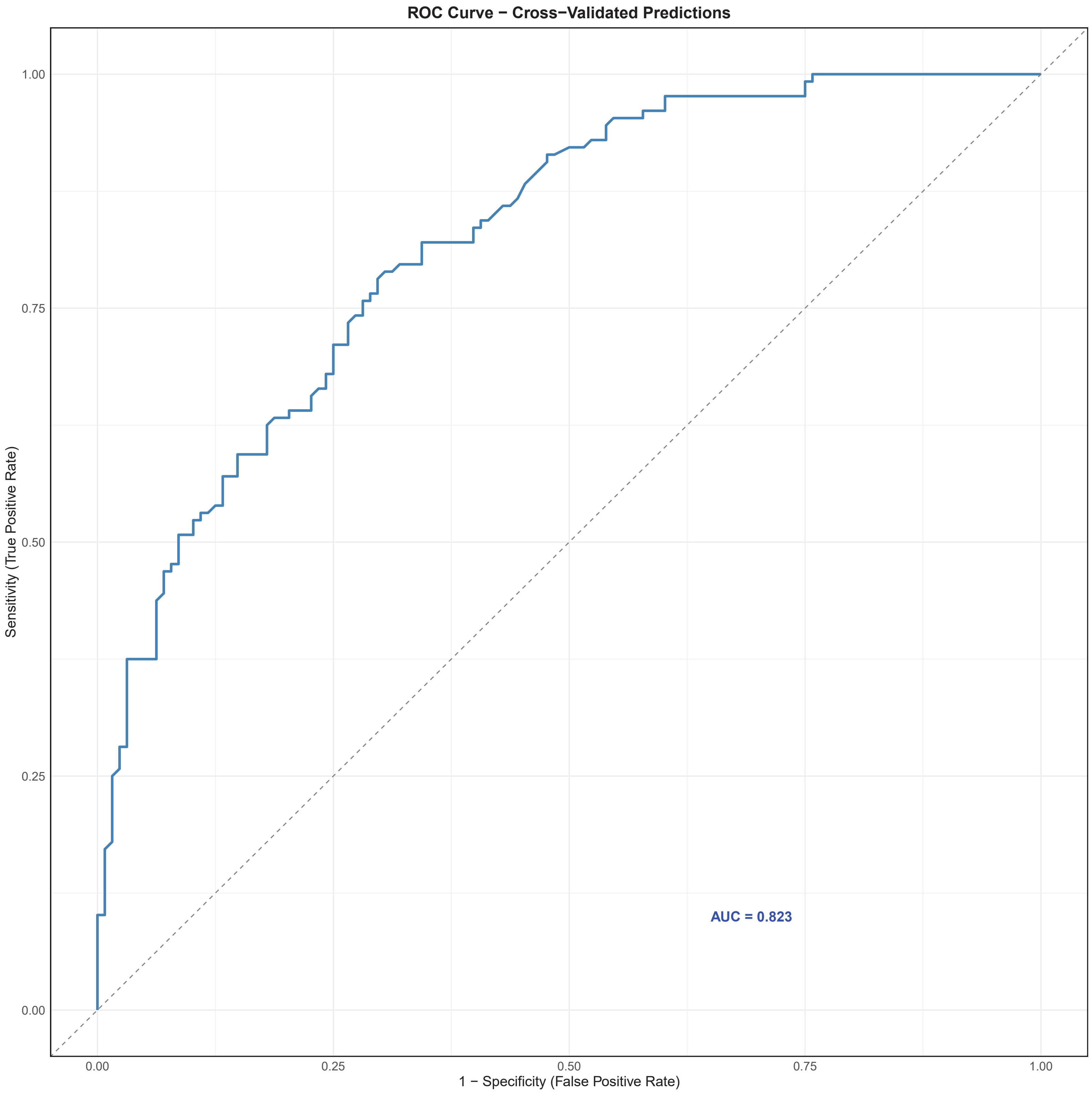

### Supplementary Figure 15

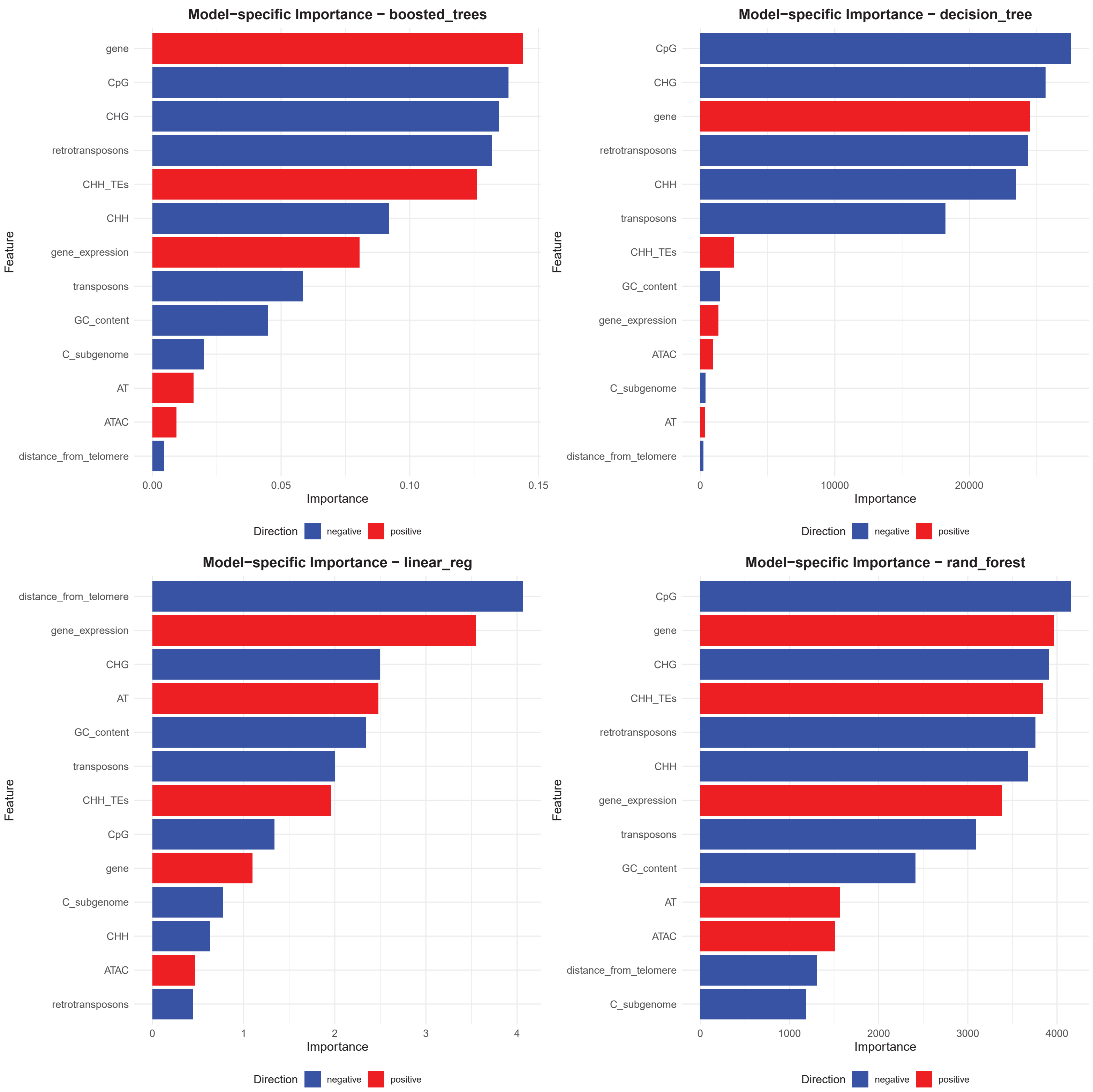

Supplementary Figure 16

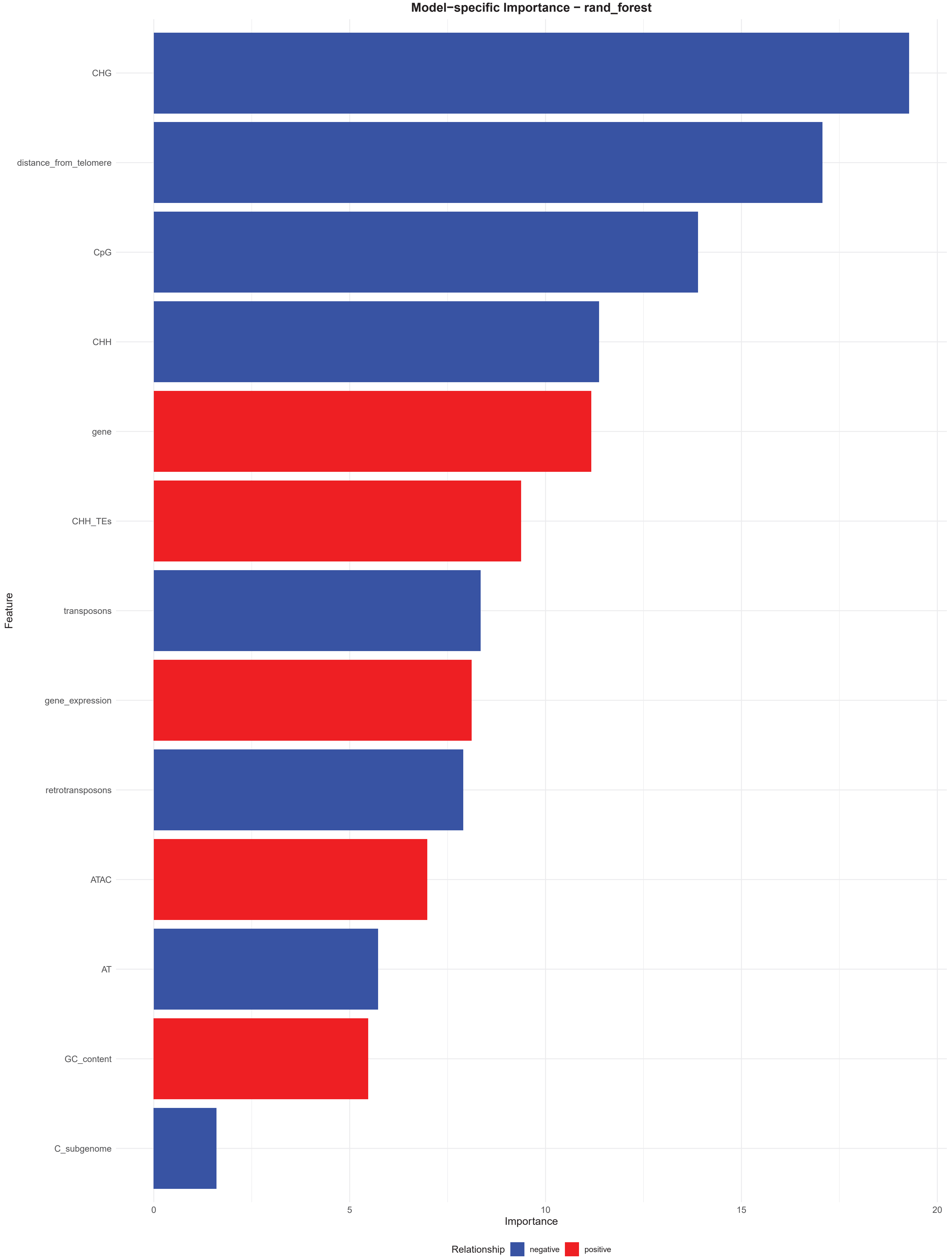

### Supplementary Figure 17

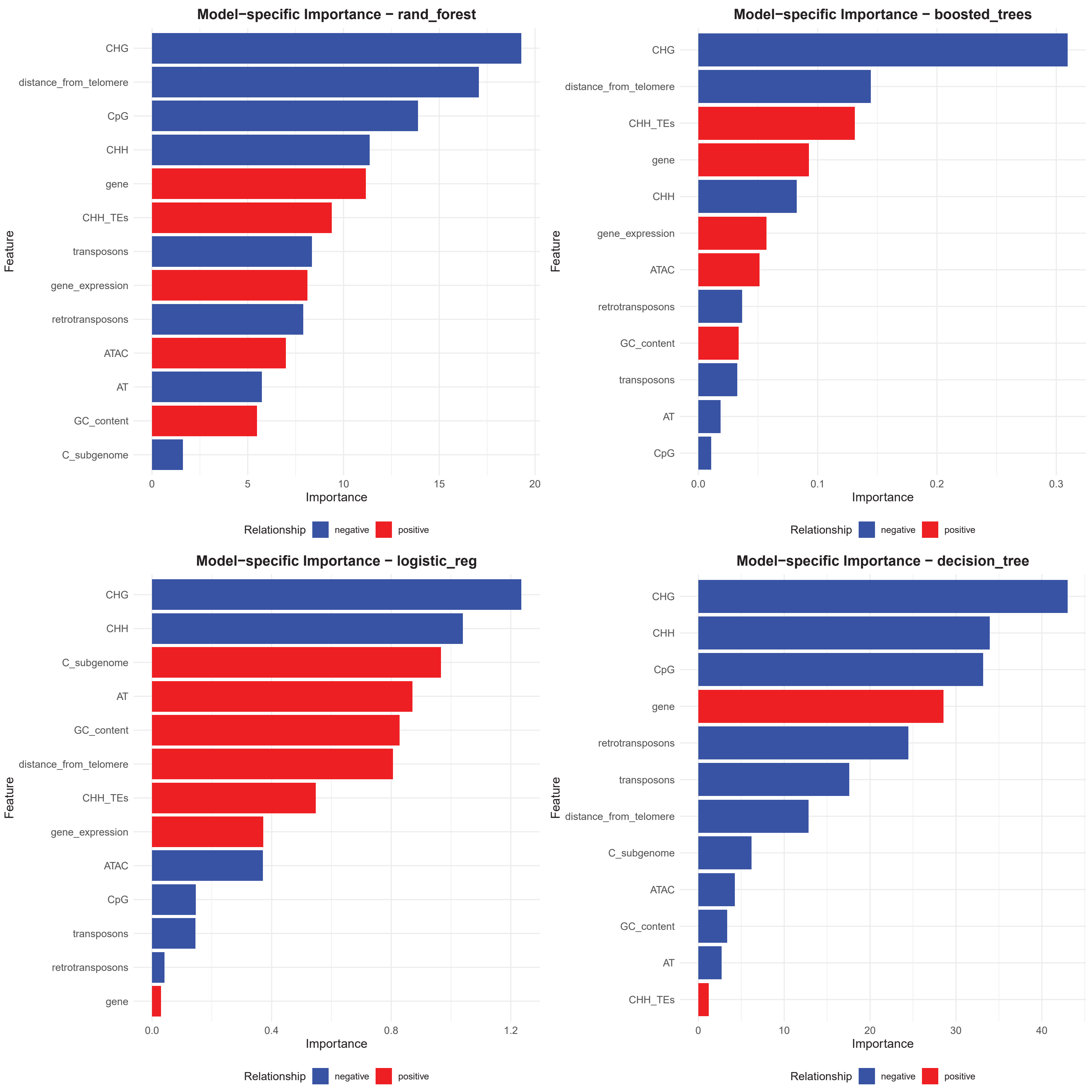

### Supplementary Figure 18

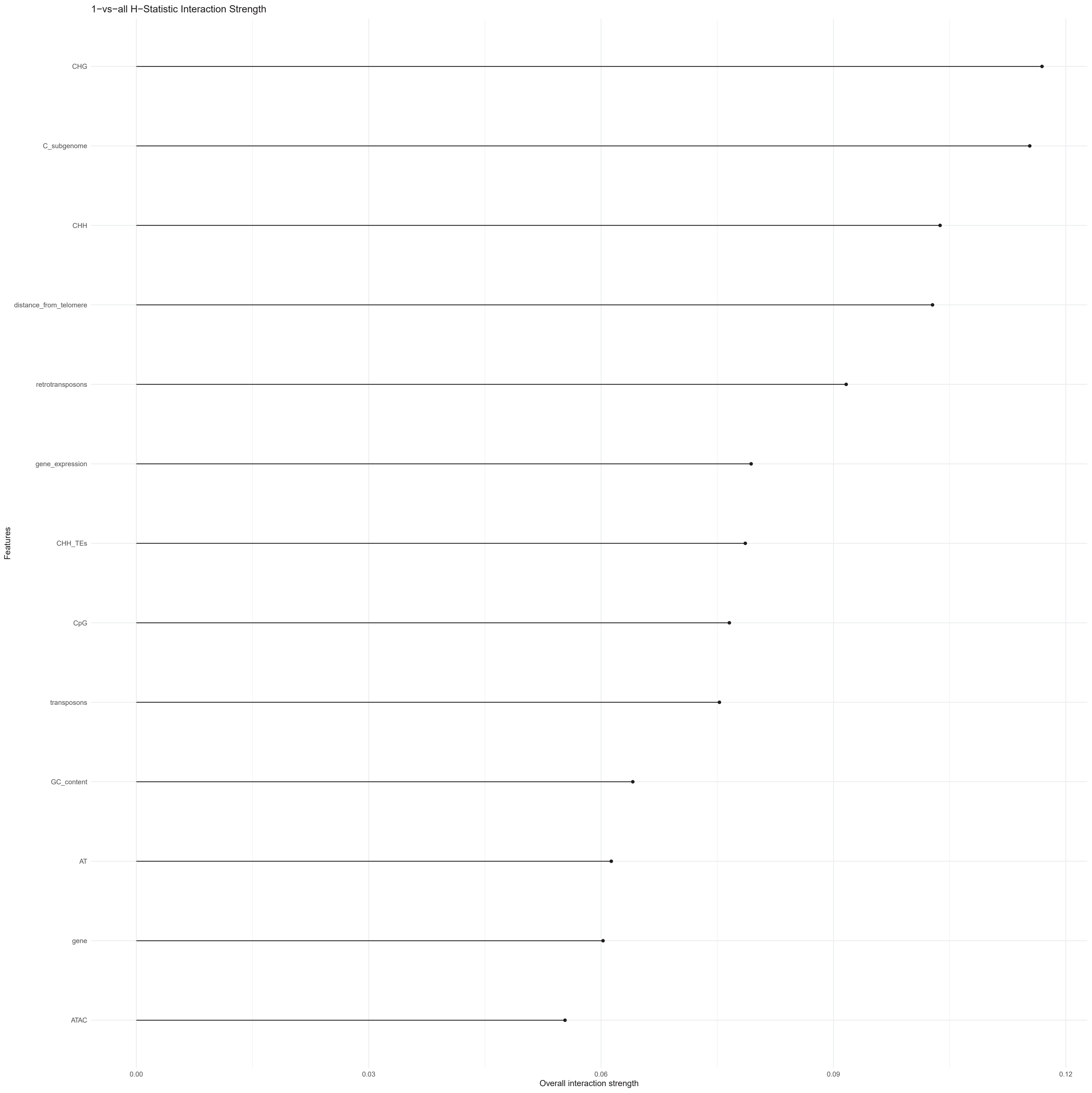

### Supplementary Figure 19

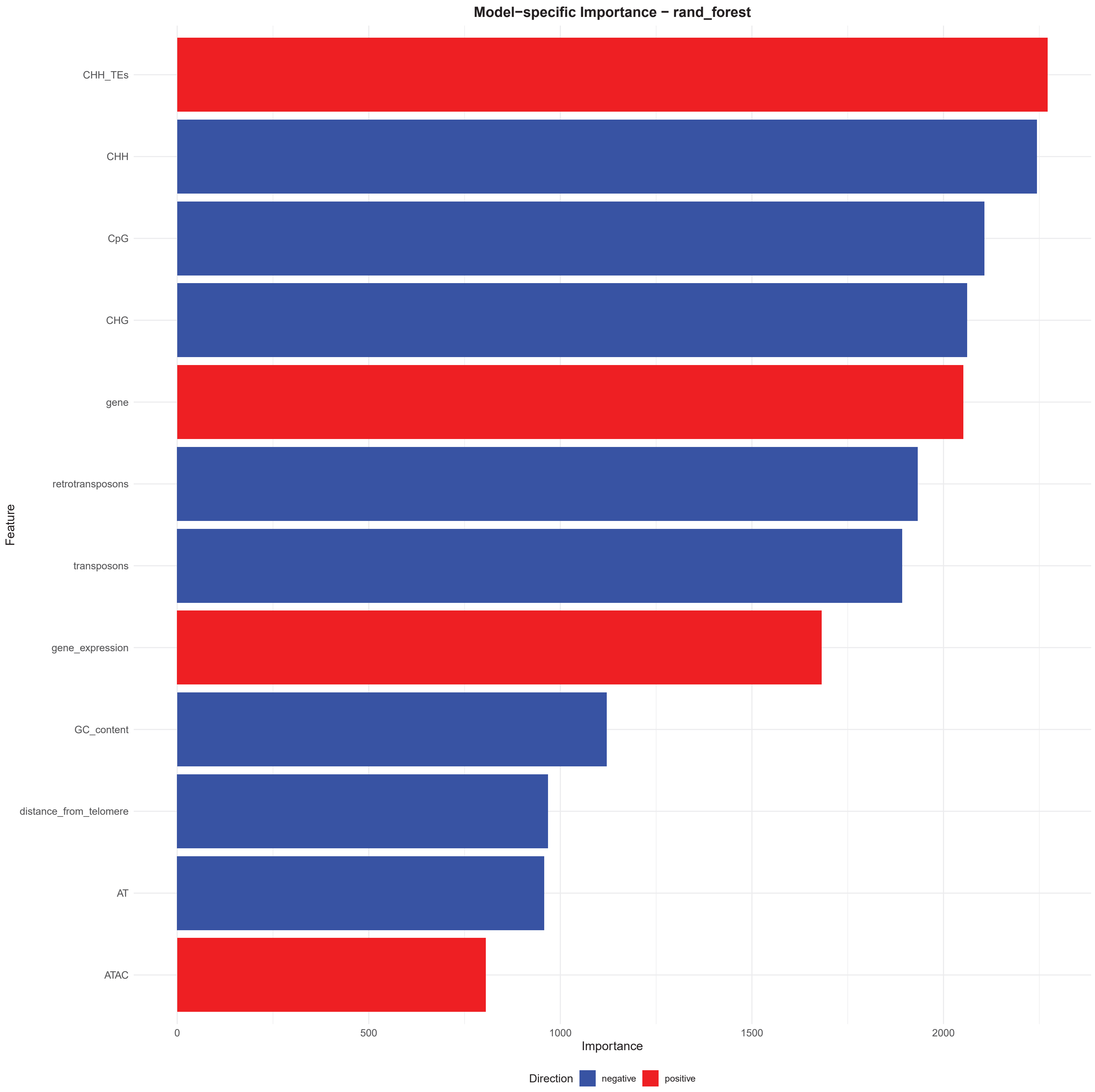

### Supplementary Figure 20

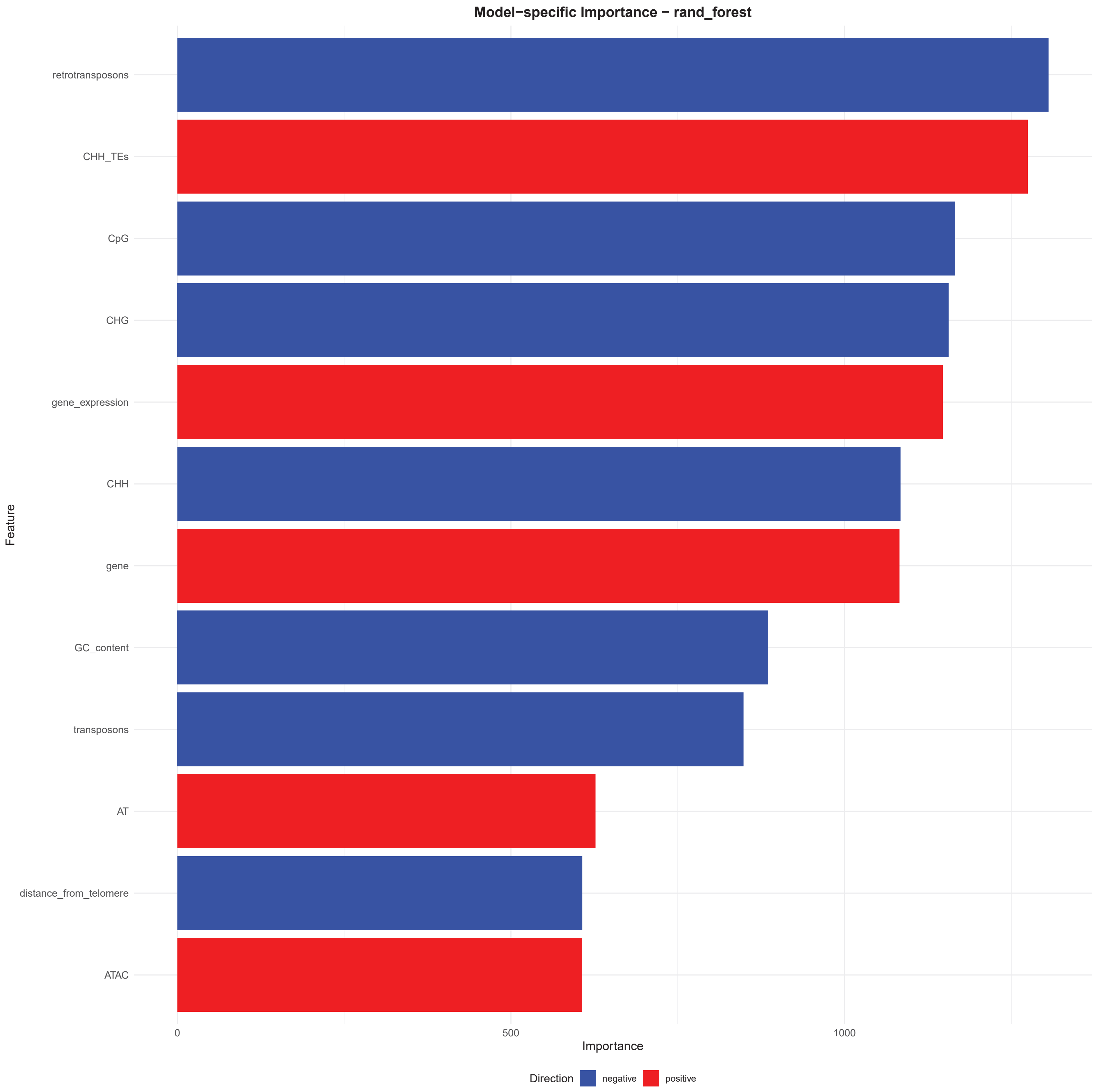

### Supplementary Figure 21

**A subgenome − 1D ALE (all features)**

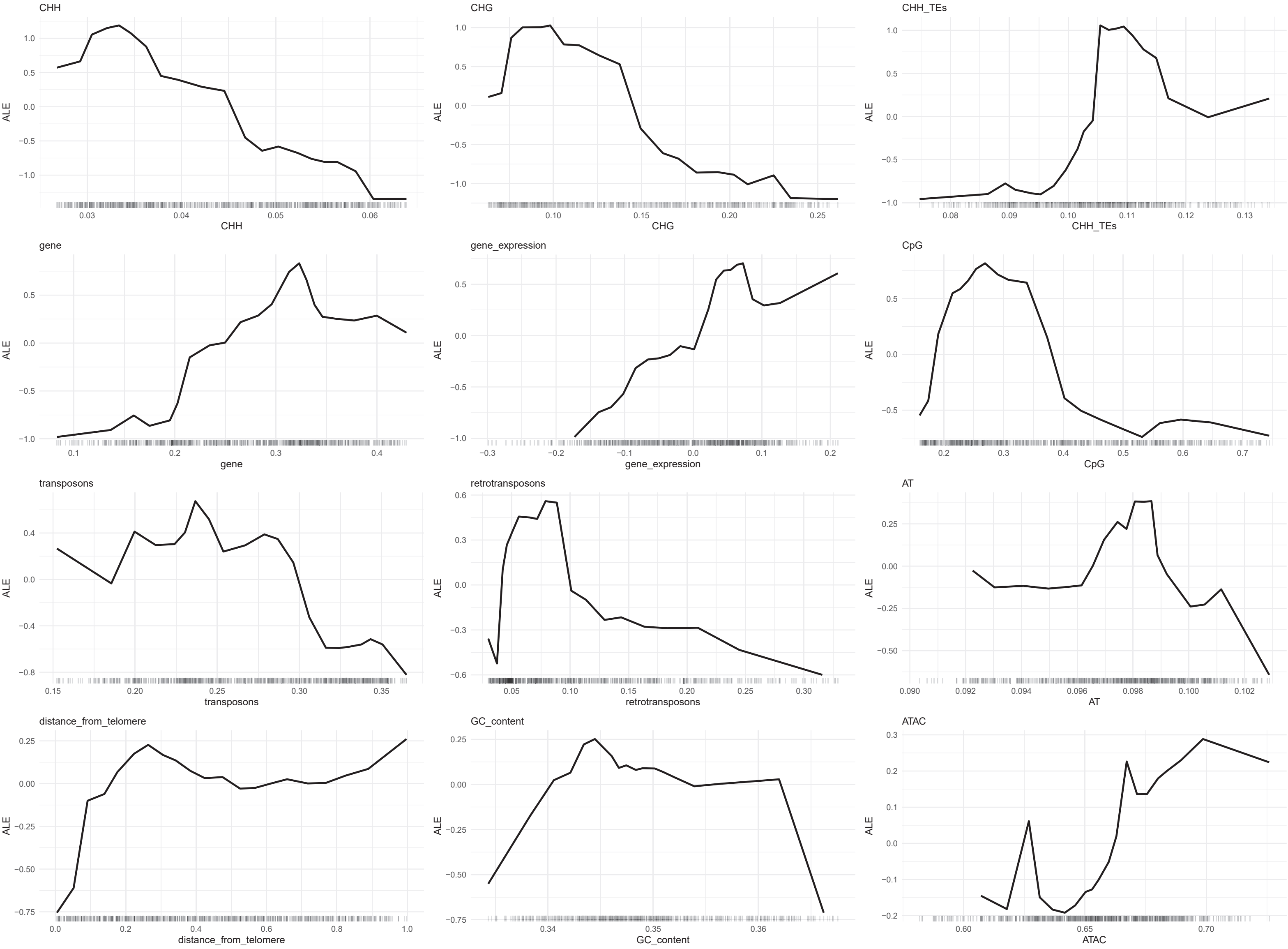

### Supplementary Figure 22

**C subgenome − 1D ALE (all features)**

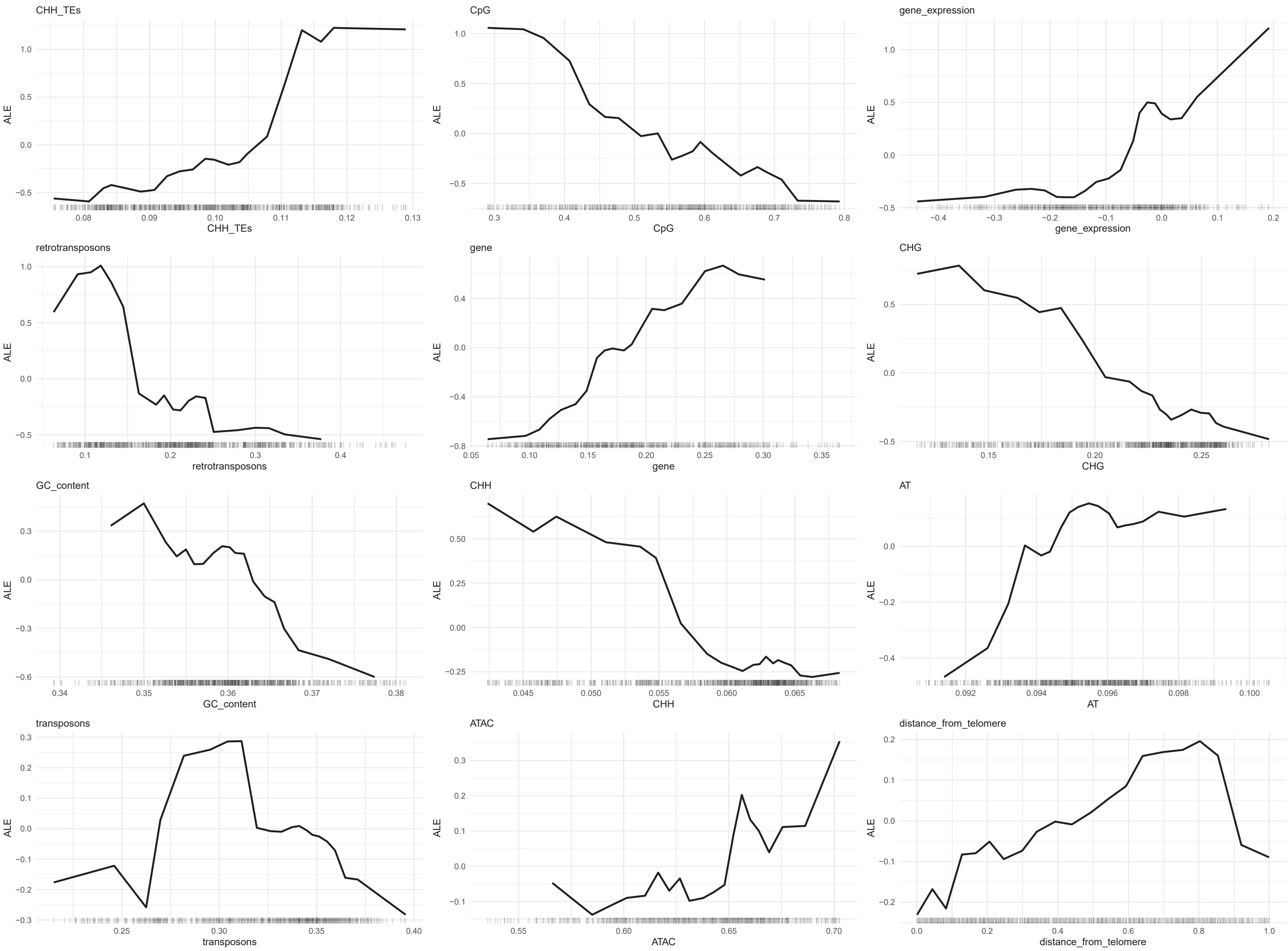
